# Targeted DNA ADP-ribosylation triggers templated repair in bacteria and base mutagenesis in eukaryotes

**DOI:** 10.1101/2024.11.17.623984

**Authors:** Constantinos Patinios, Darshana Gupta, Harris V. Bassett, Scott P. Collins, Charlotte Kamm, Anuja Kibe, Yanyan Wang, Chengsong Zhao, Katie Vollen, Christophe Toussaint, Kathryn M. Polkoff, Thuan Nguyen, Irene Calvin, Angela Migur, Ibrahim S. Al’Abri, Tatjana Achmedov, Alessandro Del Re, Antoine-Emmanuel Saliba, Nathan Crook, Anna N. Stepanova, Jose M. Alonso, Chase L. Beisel

## Abstract

Base editors create precise genomic edits by directing nucleobase deamination or removal without inducing double-stranded DNA breaks. However, a vast chemical space of other DNA modifications remains to be explored for genome editing. Here, we harness the bacterial anti-phage toxin DarT2 to append ADP-ribosyl moieties to DNA, unlocking distinct editing outcomes in bacteria versus eukaryotes. Fusing an attenuated DarT2 to a Cas9 nickase, we program site-specific ADP-ribosylation of thymines within a target DNA sequence. In tested bacteria, targeting drives efficient homologous recombination in tested bacteria, offering flexible and scar-free genome editing without base replacement nor counterselection. In tested eukaryotes including yeast, plants and human cells, targeting drives substitution of the modified thymine to adenine or a mixture of adenine and cytosine with limited insertions or deletions, offering edits inaccessible to current base editors. Altogether, our approach, called append editing, leverages the addition of a chemical moiety to DNA to expand current modalities for precision gene editing.

## INTRODUCTION

In the expanding field of genome editing, targeting chemical modifications to a specific DNA sequence offers an effective way to create precise genomic edits without relying on double-stranded DNA breaks^1–3^. These modifications are installed at selected sites by base editors (BEs) comprising an enzymatic DNA domain and a programmable DNA binding protein. After the BE acts on recognized bases within a selected target site, the modified bases then change identity, resulting in a permanent genetic substitution. As this process does not actively generate double-stranded DNA breaks at the target site, unintended and possibly harmful genetic alterations such as random insertions or deletions (indels), chromosomal abnormalities, chromothripsis are avoided^1,4^. To date, BEs have been applied in all three domains of life^5,6^ including DNA-containing organelles like mitochondria^7^, can convert each of the four bases^6^, and have recently entered clinical use^8^.

Within these advances, BEs have consistently relied on DNA deaminases to remove an amino group, changing the base’s perceived identity, or DNA glycosylases to remove the entire base, driving the base’s replacement via base excision repair^2,9^. While such “subtractive” DNA modifications represent powerful means to elicit precise gene edits, what remains unexplored is the impact of “additive” DNA modifications. Extensive work in DNA repair has shown that appended chemical moieties can elicit diverse DNA repair pathways, such as homologous recombination, translesion synthesis, nucleotide-excision repair or Fanconi anemia repair, extending well beyond base-excision repair^10–12^. However, the programmable addition of chemical moieties to DNA for gene editing remains to be explored. One promising starting point derives from the DNA ADP-ribosyltransferase protein DarT2^13^. DarT2 is part of the DarT2/DarG toxin-antitoxin system recently associated with a growing collection of anti-phage defenses (**Fig. 1a**)^14^. As the system’s toxin, DarT2 appends a single ADP-ribosyl moiety to the N3 position of thymine in single-stranded DNA using the metabolic cofactor NAD^+^ as a substrate^15^. The antitoxin DarG protein catalytically removes the appended ADP-ribosyl moiety and serves as a DNA mimic that binds DarT2^16^. During a phage infection, DarG is inactivated through an unknown mechanism, and DarT2 begins ADP-ribosylating DNA within the bacteriophage and host genome^14^. An appended ADP-ribosyl moiety interferes with DNA replication, which can block bacteriophage replication and induce cellular growth arrest. In *Escherichia coli*, growth arrest could be partially relieved through bypass via RecF-mediated homologous recombination with the sister chromatid followed by removal through nucleotide-excision repair (**Fig. 1b**)^17^. Critically, this mode of repair contrasts with traditional base editing in this bacterium^18,19^, suggesting that the installation of an ADP-ribosyl moiety could unlock distinct types of genome edits. Here, we explore such an approach, which we call append editing. As we append an ADP-ribosyl (ADPr) moiety to thymine, the approach can be abbreviated as ADPr-T append editing (ADPr-TAE).

**Fig. 1:**
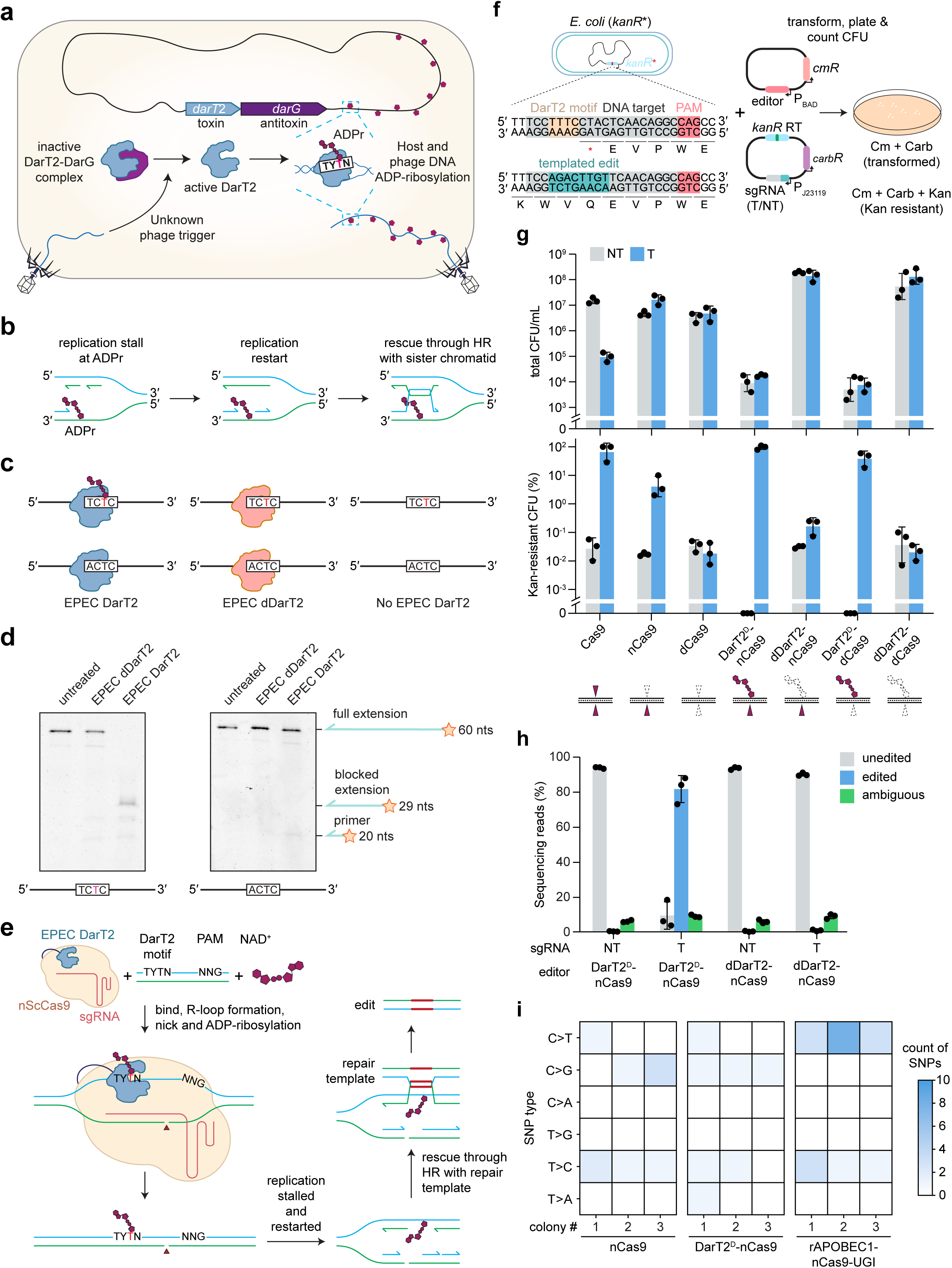
Targeted DNA ADP-ribosylation drives template-mediated homologous recombination in *E. coli*. **a,** Role of the bacterial DarT2 toxin in anti-phage immunity. **b,** Conceptualized impact and resolution of DNA ADP-ribosylation on DNA replication in *E. coli*. Based on ref. ^17^. **c,** Experimental setup for the *in vitro* polymerase-blocking assay. EPEC DarT2 recognizes the 5′-TCTC-3′ but not the 5′-ACTC-3′ motif. dDarT2: DarT2 with catalytically inactivating E170A mutation. **d,** Impact of DNA ADP-ribosylation by DarT2 on DNA polymerase extension *in vitro*. Gel images are representative of duplicate independent experiments. See Figure S1 for additional controls. **e,** Configuration of the append editor utilizing DarT2. The editor combines ScCas9 mutated to nick the target DNA strand and a fused DarT2 that ADP-ribosylates the non-target DNA strand displaced as part of R-loop formation. This combination is predicted to drive homologous recombination with a provided repair template. **f,** Experimental setup for reverting a prematurely terminated kanamycin resistance gene (*kanR*\*) in *E. coli*. The chromosomally integrated gene contains a premature stop codon that is reverted as part of homologous recombination, thus conferring kanamycin resistance. RT: DNA repair template. **g,** Impact of programmable DNA ADP-ribosylation on cell viability and kanamycin-resistance frequency. Bars and error bars represent the geometric mean and geometric s.d. of three independent experiments started from separate transformations. Dots represent individual measurements. CFU: colony-forming units. T sgRNA: sgRNA with a guide targeting the intended site. NT sgRNA: sgRNA with a non-targeting guide. Below: cartoons designate whether a given DNA strand is unaltered, nicked or ADP-ribosylated. **h,** Amplicon sequencing of the *kanR\** target site from batch cultures. Bars and error bars represent the mean and s.d. of three independent experiments starting from separate transformations. Dots represent individual measurements. **i**, Genome-wide profiling of off-target edits. The indicated editor was expressed with a non-targeting sgRNA and in the absence of an RT. See Table S1 for more information on the identified edits. Whole-genome sequencing was performed on genomic DNA extracted from cultures beginning with an individual colony. Both strands are considered for a given edit (e.g., T > A and A > T are combined).

## RESULTS

### CRISPR-guided ADP-ribosylation drives homologous recombination in *E. coli*

To explore the outcome of targeted DNA ADP-ribosylation, we selected the previously characterized DarT2 from enteropathogenic *Escherichia coli* (EPEC) O127:H6 str. E2348/69^17^. The EPEC DarT2 was shown to ADP-ribosylate single-stranded DNA at the third position in a 5′-TYTN-3′ motif (Y = C/T), with the fourth position biased against a G^17^. Paralleling its growth-inhibitory effects *in vivo*, this DarT2 blocked extension by the large fragment of *E. coli*’s DNA Polymerase I *in vitro* from a single-stranded (ss)DNA template with the recognition motif (5′-TCTC-3′), whereas extension was unhindered with a mutated motif (5′-ACTC-3′) or with DarT2 containing the inactivating E170A mutation (dDarT2) (**Figs. 1c-d and S1**)^17^.

To direct DNA ADP-ribosylation, we fused DarT2 to the N-terminus of the PAM-flexible (5′-NNG-3′) *Streptococcus canis* Cas9 (ScCas9) (**Fig. 1e**)^20^. Directing the DarT2-Cas9 fusion to a target sequence through a designed single-guide (sg) RNA would localize DarT2 to the non-target strand displaced during R-loop formation (**Fig. 1e**). If the non-target strand contains a 5′-TYTN-3′ motif accessible to DarT2, then the target thymine within the motif would be ADP-ribosylated and serve as a block to DNA replication. As wild-type DarT2 would arrest cell growth through genome-wide ADP-ribosylation, we included a previously reported spontaneous G49D mutation in the NAD^+^-binding loop helix (DarT2^D^) exhibiting reduced cytotoxicity^17^. To promote repair through a provided DNA template rather than the sister chromatid, we used a nickase version of Cas9 (D10A) that only cleaves the target strand and provided a plasmid-encoded repair template with ∼500-bp homology arms flanking the intended edits.

As a simple readout of homologous recombination, we introduced a premature stop codon into a chromosomally integrated kanamycin resistance gene in *E. coli* strain MG1655 (**Fig. 1f**). The premature stop codon overlaps with an ScCas9 target containing the 5′-TTTC-3′ DarT2 motif and a PAM sequence, while a provided repair template with ∼500-bp homology arms introduces mutations that revert the premature stop codon and remove the DarT2 motif. As part of an editing assay, plasmids encoding the editor, sgRNA, and repair template are transformed into *E. coli*, and colony counts are compared following editor induction and plating with or without kanamycin.

To set a baseline, we applied dsDNA cleavage with Cas9, which is commonly used for genome editing in bacteria^21^. As dsDNA cleavage principally removes cells that did not undergo recombination, using Cas9 resulted in 66% kanamycin-resistant colonies and a 159-fold colony reduction compared to the non-targeting control (*p* = 0.0002, n = 3) (**Fig. 1g**). The nickase version of Cas9 did not deplete colony counts (3.6-fold increase relative to the non-targeting control, *p* = 0.02, n = 3) but at the expense of fewer kanamycin-resistant colonies (4%), in line with nicking being less cytotoxic but a poor driver of homologous recombination. Binding DNA alone with a catalytically dead Cas9 (dCas9) exhibited similar colony counts to nCas9 (*p* = 0.07, n = 3) and did not drive any measurable editing.

Turning to append editing with DarT2, the nCas9-DarT2^D^ fusion yielded 96% kanamycin-resistant colonies, and negligible depletion in colony counts compared to its non-targeting control (2.0-fold increase, *p* = 0.25, n = 3) (**Fig. 1g**). Both DNA ADP-ribosylation and opposite-strand nicking were important, as conferring kanamycin resistance was less effective with nicking alone (nCas9-dDarT, 0.16%, *p* = 0.003, n = 3) or ADP-ribosylation alone (dCas9-DarT2^D^, 38%, *p* = 0.029, n = 3) when compared to nCas9-DarT2^D^. All screened kanamycin-resistant colonies contained the intended edit (**Fig. S2**). Dart2^D^ still conferred cytotoxicity, as cell counts were low even for the non-targeting controls and increased upon deactivation of DarT2 (**Fig. 1g**), creating an opportunity to further attenuate the toxin. Collectively, append editing with DarT2 drives homologous recombination with a provided template in *E. coli*, yielded editing that outperformed traditional Cas9-based approaches.

### Targeted ADP-ribosylation does not induce detectable base edits in *E. coli*

Our employed reporter assay requires homologous recombination to confer kanamycin resistance. However, chemically modifying DNA bases can lead to single nucleotide edits as demonstrated by BEs^18,22^. We therefore asked whether append editing could drive editing without antibiotic selection but also induce base mutagenesis. First, we repeated the *kanR* reporter assay in the absence of kanamycin selection and performed amplicon sequencing on the target site from liquid culture (**Fig. 1h**). Under targeting conditions, append editing yielded 82% of total reads with the desired edit that drastically dropped with nicking alone (0.9%), paralleling the fraction of kanamycin-resistant colonies (**Fig. 1g**). Of the remaining reads, the few detected substitutions of the ADP-ribosylated thymine were not significantly elevated in any sample (*F* = 1.03, *p* = 0.39, *df* = 3) (**Fig. S3**). As homologous recombination could overshadow base editing, we performed the assay in the absence of the repair template. However, the 16 screened colonies only yielded the original sequence (**Fig. S4**). Therefore, append editing with DarT2 did not result in detectable base edits in *E. coli*, further supporting sole triggering of homologous recombination.

Base editing can also occur at genomic sites unrelated to the target sequence presumably through the DNA modification domain acting on DNA that is temporarily single-stranded ^23^. Given the lack of obvious substitutions at the target site with append editing, we hypothesized that DarT2 expression would not lead to such edits associated with BEs. Culturing editor-expressing cells and performing whole-genome sequencing of three individual clones (**Fig. 1i and Table S1**), a cytosine base editor (CBE) yielded the expected C-to-T edits^23^, with either three or eight edits in each clone. In contrast, the ADPr-TA editor yielded no T-to-G edits and similarly few T-to-C edits as the CBE or no editor. One of the three clones with the ADPr-TA editor yielded a single T-to-A edit, while none were observed with the CBE or no editor. This one edit was associated with the 5′-TYTN-3′ motif, suggesting that base mutagenesis is possible but rare (**Table S1**). Thus, even a highly active DarT2 that reduces cell viability (**Fig. 1g**) does not inherently drive base edits across the *E. coli* genome.

### Attenuating DarT2 alleviates cytotoxicity without compromising homologous recombination

ADPr-TAE yielded high editing efficiencies, although the expressed DarT2^D^ exhibited strong cytotoxicity (**Fig. 1g**). As the cytotoxicity was likely due to ADP-ribosylation of ssDNA across the genome, we aimed to attenuate DarT2 without compromising localized ADP-ribosylation and subsequent initiation of homologous recombination using structural insights and sequence conservation (**Fig. 2a**). While the structure of EPEC DarT2 remains to be experimentally determined, a crystal structure is available for the *Thermus* sp. 2.9 DarT2 sharing 34% amino-acid identity with EPEC DarT2^15^. Aligning this structure with the AlphaFold-predicted structure of EPEC DarT2^24^, we selected a subset of residues potentially involved in binding the DNA recognition motif (M84, M86, R57, R92, R166) or potentially flanking regions of the DNA strand not captured in the crystal structure (R193). The positively charged arginines were mutated to uncharged alanine, while the methionines were mutated to leucine to disrupt the coordinating sulfur while preserving the residue’s hydrophobicity and chain length. Testing these substitutions in combination with G49D as part of the kanamycin-resistance reversion assay (**Figs. 1f**), we found that all improved cell viability (**Fig. 2b**). At the same time, three of the mutations (M86L, R92A, R193A) maintained the fraction of kanamycin resistant colonies comparable to the original G49D (*p* = 0.77, 0.51 and 0.27 respectively, n = 3) (**Fig. 2b**), representing candidates for further use with append editing.

**Fig. 2:**
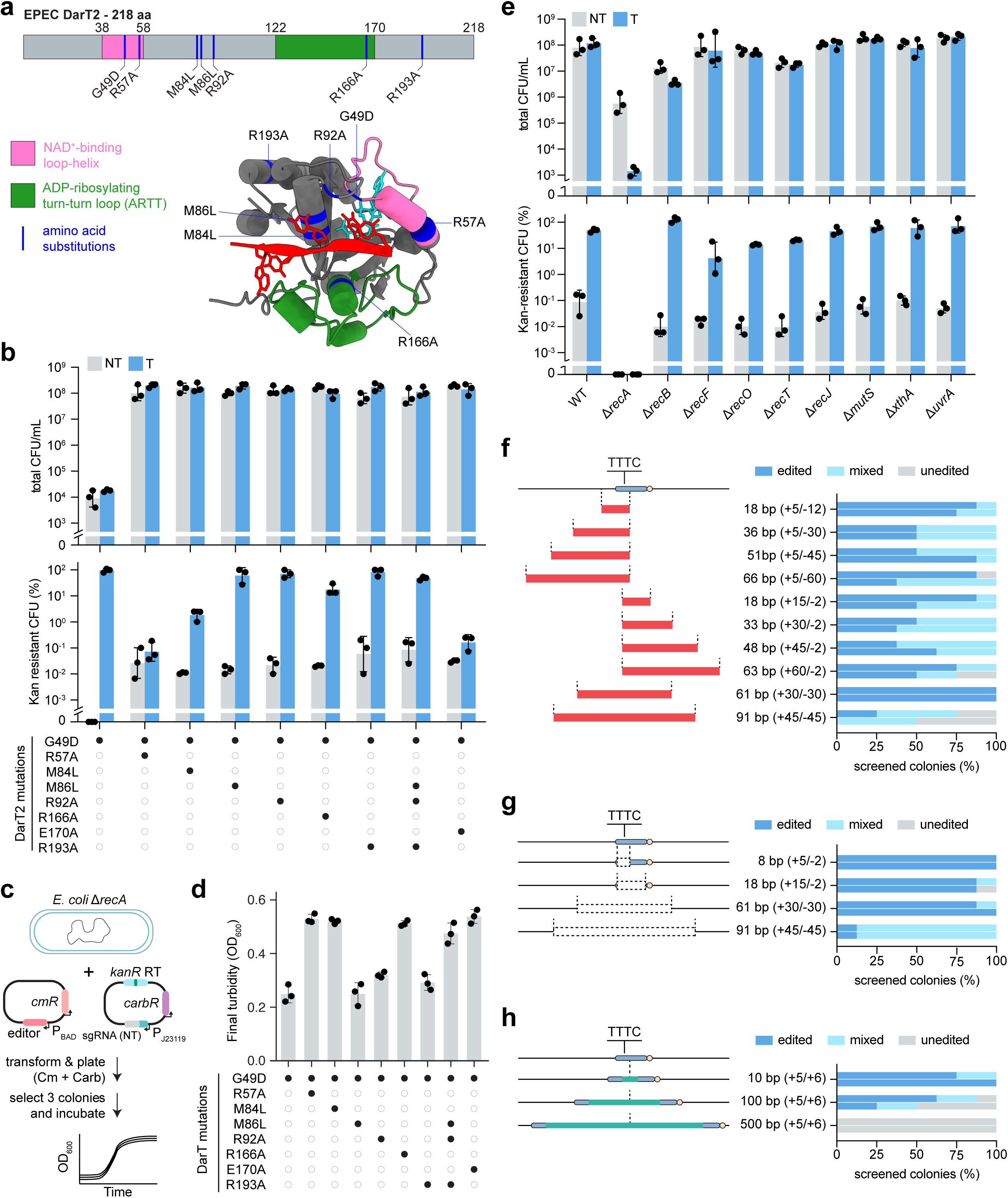
Attenuating DarT2 alleviates cytotoxicity while mediating efficient and flexible gene editing in *E. coli*. **a,** Predicted structure of EPEC DarT2. Tested substitutions are in blue. **b,** Impact of tested substitutions on cell viability and kanamycin-resistance frequency. See Figure 1f for the experimental setup. **c,** Experimental setup for assessing growth defects caused by editor expression in a Δ*recA*-deletion strain of *E. coli*. **d,** Impact of expressing an append editor with the indicated DarT2 mutant with a non-targeting sgRNA in the Δ*recA*-deletion strain of *E. coli*. Endpoint optical density (OD) measurements were taken after 12 h of culturing. See Figure S5 for growth curves. Bars and error bars in b and e represent mean ± geometric s.d. of three independent experiments started from separate transformations. Dots represent individual measurements. **e,** Impact of deleting DNA repair genes on cell viability and kanamycin-resistance frequency. Bars and error bars in b and e represent geometric mean ± geometric s.d. of three independent experiments starting from separate transformations. Dots represent individual measurements. **f,** Introducing sequence replacements with ADPr-TA editing. **g,** Introducing deletions with ADPr-TA editing. **h,** Introducing insertions with ADPr-TA editing. For f-h, Left: size and location of substitutions (red bar), deletions (dashed box), or insertions (green bar). Numbers (e.g., +5/-12) indicate the edited region in relation to the ADP-ribosylated thymine. Right: fraction of screened colonies containing the intended edit. Each bar represents one of two biological replicates starting from separate transformations, screening at least 8 colonies per biological replicate. See Figure S7 for examples of Sanger sequencing chromatograms indicating edited, mixed, and unedited colonies.

Viability was greatly enhanced across the single-substitution variants, yet DarT2 may still exert target-independent ADP-ribosylation that could have more subtle effects on cell growth and behavior. We therefore generated cells hypersensitive to ADP-ribosylation by deleting the core repair gene *recA* to disable homologous recombination, and we assessed cell growth when expressing each ADPr-TAE variant under non-targeting conditions (**Figs. 2c and S5**). While growth rates in exponential phase were similar (**Fig. S5**), we observed marked differences in entry into stationary phase. In particular, amino acid substitutions that previously compromised editing (M84L, R57A, R166A) yielded final turbidities paralleling the inactivating E170A (*p* = 0.35, 0.65, and 0.22 respectively, n = 3) (**Figs. 2d and S5**). In contrast, substitutions that previously showed high editing efficiencies (M86L, R92A, R193A) exhibited a final turbidity similar to G49D alone (*p* = 0.99, 0.05, and 0.17 respectively, n = 3) and lower than the E170A. We therefore combined the high editing mutations (M86L, R92A, R193A) into a four-substitution version of DarT2, DarT2^DLAA^. This version maintained cell viability and a high frequency of kanamycin-resistant colonies (49%) in *E. coli* MG1655 (**Fig. 2b**). Moreover, in the *recA*-deletion strain, the append editor with DarT2^DLAA^ restored final turbidity to approach that of the editor lacking ADP-ribosylation (*p* = 0.09, n = 3) (E170A) (**Fig. 2d**).

By improving cell viability and growth in a strain in which homologous recombination was fully disabled, the append editor with DarT2^DLAA^ afforded the opportunity to probe the genetic basis of templated-mediated editing. Prior work on the cytotoxicity of DarT2^D^ in *E. coli* revealed a key role by RecF and possibly nucleotide-excision repair^17^. However, the involved DNA repair pathways as part of targeted ADP-ribosylation with opposite-strand nicking could differ. Within the kanamycin-reversion assay (**Fig. 1f**), *recA* was essential for editing and even showed some reduction in colony counts under non-targeting conditions (**Fig. 2e**). Disrupting the RecBCD branch of recombination (Δ*recB*) reduced viability but also increased the frequency of kanamycin-resistant colonies, suggesting a role in survival in the absence of recombination with the provided repair template. In contrast, disrupting the alternative RecFOR recombination pathway (Δ*recF*, Δ*recO*) reduced editing relative to the wild type (one-sided Welch’s t-test, *p* = 0.048, 0.001 respectively, n = 3) but not viability for *recF* (one-sided Welch’s t-test, *p* = 0.40, n = 3), suggesting involvement in templated recombination. Disrupting RecA-independent RecT recombination (Δ*recT*) significantly reduced both viability and editing (one-sided Welch’s t-test, *p* = 0.002, 0.003 respectively, n = 3), suggesting involvement in both survival and templated recombination. Finally, the DNA repair exonuclease RecJ (Δ*recJ*), mismatch repair (Δ*mutS*), base excision repair (Δ*xthA*) and nucleotide-excision repair (Δ*uvrA*) did not impact editing (one-sided Welch’s t-test, *p* = 0.89, 0.68, 0.81 respectively, n = 3) or viability (one-sided Welch’s t-test, *p* = 0.87, 0.24, 0.93, n = 3) relative to the wild type. These findings implicate multiple recombination pathways as part of ADPr-TAE *in E. coli*.

### Attenuated ADP-ribosylation enables flexible and non-cytotoxic genome editing in bacteria

Append editing with DarT2^DLAA^ efficiently reverted the premature stop codon in the kanamycin-reversion assay. However, the reliance on homologous recombination lends to a much broader range of edits in different genes and bacteria. We therefore explored the bounds of ADPr-TA editing. For simplicity, editing was performed around the premature stop codon in the kanamycin-reversion assay. When testing edits beyond reversion of the stop codon, editing efficiency was determined without kanamycin selection by assessing the target-site size or sequence of individual colonies.

Beginning with the homology arms, condensing their length from ∼500 to 100 bp reduced the frequency of kanamycin resistance from 86% to 28%, while arm lengths of 50 bp and below exhibited virtually no kanamycin resistance (**Fig. S6**). Continuing with ∼500-bp homology arms, we tested increasingly larger replacements, deletions and insertions (**Fig. 2f-h**). Replacements extending up to 60 bp upstream or downstream of the target site or 91 bp spanning the target site were present in 80-100% and 50-75% of screened colonies, respectively, either as complete or partial conversions (**Figs. 2g and S7**). Separately, deletions up to 91 bp were present in 90-100% of screened colonies, albeit with a high fraction of partial conversion with the largest deletion. Finally, insertions of 10 bp and 100 bp were present in 100% and 50-90% of screened colonies, respectively. No colonies contained an insertion of 500 bp (**Fig. S8**), indicating an upper limit to recombination. Editing was not limited to this target site in *E. coli*, as we could introduce substitutions at four additional targeted genes in *E. coli* (**Fig. S9a**) as well as one targeted gene in the pathogen *Salmonella enterica* (**Fig. S9b**). Collectively, ADPr-TAE can introduce ranging replacements, insertions, and deletions in bacteria without sacrificing viability.

### Targeted ADP-ribosylation preferentially drives base mutagenesis in yeast and plants

Given that append editing drove templated recombination in bacteria, we asked whether eukaryotes would undergo similar editing outcomes. Beginning with baker’s yeast *Saccharomyces cerevisiae* cultured as haploids, we transformed plasmids encoding the DarT2^DLAA^ append editor, an sgRNA and a repair template with ∼250-bp homology arms to introduce a premature stop codon as part of six substitutions in the *FCY1* gene. Individual colonies were then screened based on Sanger sequencing of the target site (**Figs. 3a and S10**). This approach resulted in 17% of screened colonies containing the templated substitution (**Fig. 3b**). No edited colonies were obtained under non-targeting conditions or with DNA nicking alone, affirming the necessity of targeted ADP-ribosylation.

**Fig. 3:**
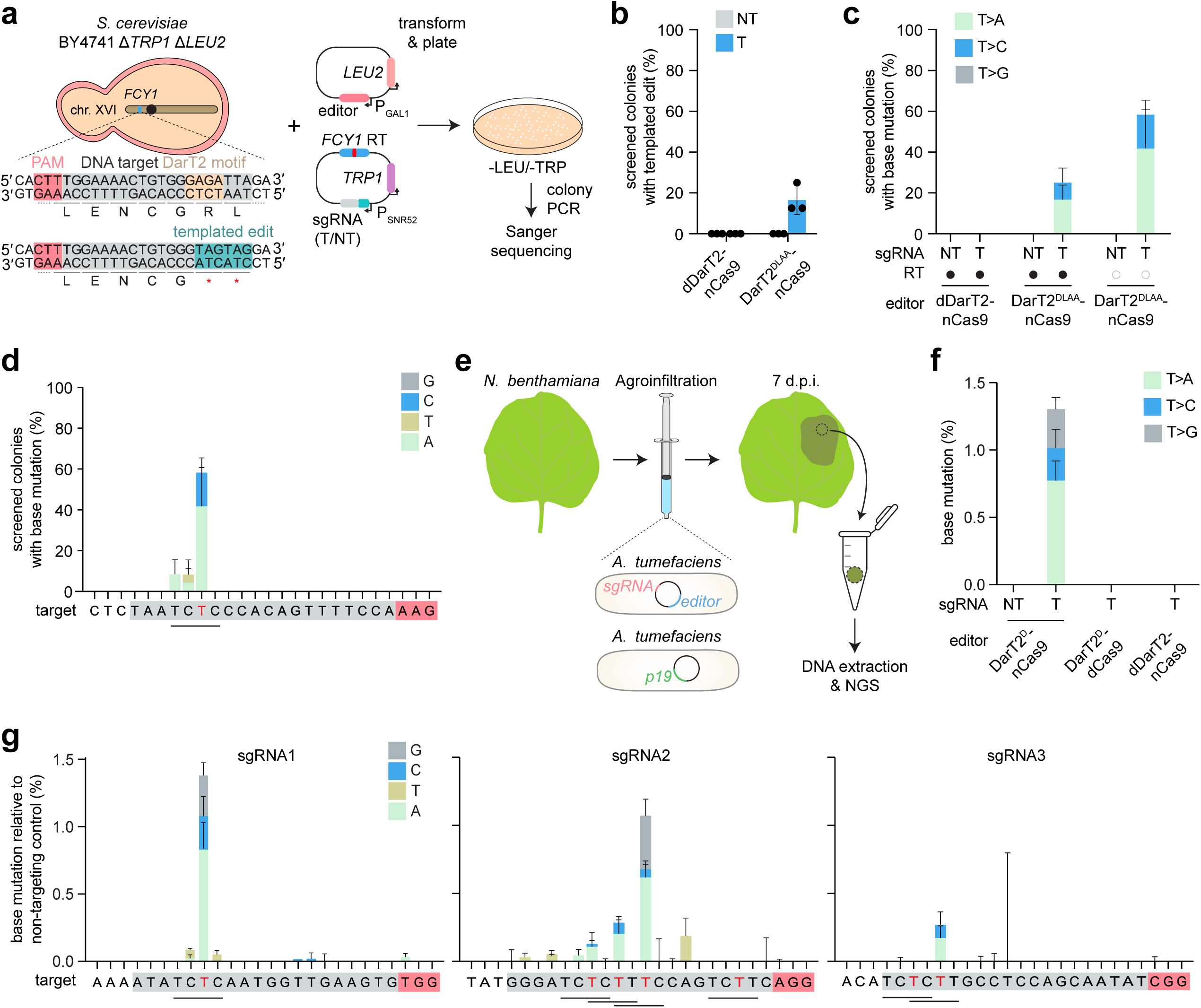
Programmable DNA ADP-ribosylation primarily drives base mutagenesis in yeast and plants. **a,** Experimental setup for introducing a six-base replacement with two adjacent premature stop codons in the *FCY1* gene of *S. cerevisiae*. **b,** Impact of ADPr-TAE on templated recombination in the presence of a RT. Bars and error bars represent the mean and s.d. of three independent experiments started from separate transformations. Dots represent individual measurements. **c,** Impact of ADPr-TAE on mutagenesis of the ADP-ribosylated thymine in the presence or absence of a RT. **d,** Frequency of base mutations across the sgRNA target. Each black bar specifies DarT2 recognition motifs, while the red base specifies the ADP-ribosylated base within the motif. See Figure S10-S11 for representative Sanger sequencing chromatograms. For c-d, bars and error bars represent the mean and =s.d. of three independent experiments started from separate transformations. **e,** Experimental setup for assessing ADPr-TAE without a repair template in *N. benthamiana*. **f,** Frequency of base mutagenesis of the ADP-ribosylated thymine in the sgRNA 1 target in the *PDS1* gene compared to the non-targeting control. **g,** Frequency of base mutations across the DNA target for sgRNA1-3 compared to the non-targeting control. See Figure S13 for the location of base mutations under targeting and non-targeting conditions. For f-g, bars and error bars represent the mean and=SEM of three independent experiments started from separate transformations.

Beyond templated recombination, we observed a distinct set of edits in 25% of screened colonies: conversion of the ADP-ribosylated thymine into a different base (**Figs. 3c and S10**). These base substitutions principally occurred at the thymine expected to undergo ADP-ribosylation by DarT2, with the modified base becoming an A (67%) or a C (33%) (**Fig. 3c**). Homologous recombination and base mutagenesis represented mutually exclusive repair outcomes, as removing the repair template enhanced the mutagenesis frequency without altering the location and distribution of mutations (**Figs. 3c-d and S11**). Base mutation was also observed when targeting sites within the genes *ALP1* and *JSN1*, albeit at lower frequencies (**Fig. S12**). Thus, in yeast, append editing drives either homology-directed repair or mutagenesis of the ADP-ribosylated thymine.

The outcomes of append editing in yeast represented a major deviation from what we observed in tested bacteria and could reflect distinct editing outcomes in eukaryotes at large. However, in contrast to higher eukaryotes, yeast engages in non-homologous end joining less frequently and lacks poly-ADP-ribosyl polymerases involved in dsDNA break repair that add and extend ADP-ribosyl groups on DNA ends^25,26^. We therefore assessed the impact of ADPr-TAE in the model plant *Nicotiana benthamiana*. As a simple and fast assay, *Agrobacterium* constructs encoding the append editor are injected into *N. benthamiana* leaves, and the type and frequency of edits are assessed via target amplicon sequencing from transfected tissues (**Fig. 3e**). In this setup, no repair template was included given the generally low frequencies of homologous recombination in this type of transfection assay in plants^27^. We also used the *Streptococcus pyogenes* Cas9 (SpCas9) given the availability of existing constructs, and we fused DarT2^D^, which did not result in any obvious morphological changes.

Despite expectedly low transfection efficiencies, we could measure substitution of the ADP-ribosylated thymine as the dominant outcome in 1.4% of reads targeting the *PDS1* gene (**Fig. 3f-g**). The thymine was converted to the other three bases, but with a bias toward A (59%) over C (19%) and G (22%). Testing two other target sites within *PDS1*, including one containing multiple DarT2 motifs, resulted in similar mutagenesis of the ADP-ribosylated T, with a bias toward A (**Figs. 3g and S13**). Indels were observed in targeting samples, but at frequencies 6-80-fold lower than base mutagenesis (**Fig. S14**). Thus, append editing can drive mutagenesis of the ADP-ribosylated base in both yeast and plants, reflecting distinct editing outcomes from those we observed in bacteria.

### Targeted ADP-ribosylation drives base mutagenesis in human cells lacking TARG1

As a final but important branch of eukaryotes, we sought to explore append editing in human cells. Unlike yeast and plants, human cells possess an ADP-ribosyl deacylase TARG1 that was previously shown to reversibly remove the ADP-ribosyl moiety appended to thymines by DarT2 (**Fig. 4a**)^28^. We therefore began by assessing ADPr-TAE in human cells with an intact or disrupted *TARG1* gene (**Fig. S15**). Plasmid constructs encoding an SpCas9-based editor and an sgRNA were transiently transfected into HEK293T cells, and editing was assessed through next-generation sequencing of the target site in *EMX1* without sorting or selection of transfected cells (**Fig. 4b**). An oligonucleotide repair template specifying a nine-base substitution and four-base deletion was included to evaluate both homologous recombination and base mutagenesis in parallel.

**Fig. 4:**
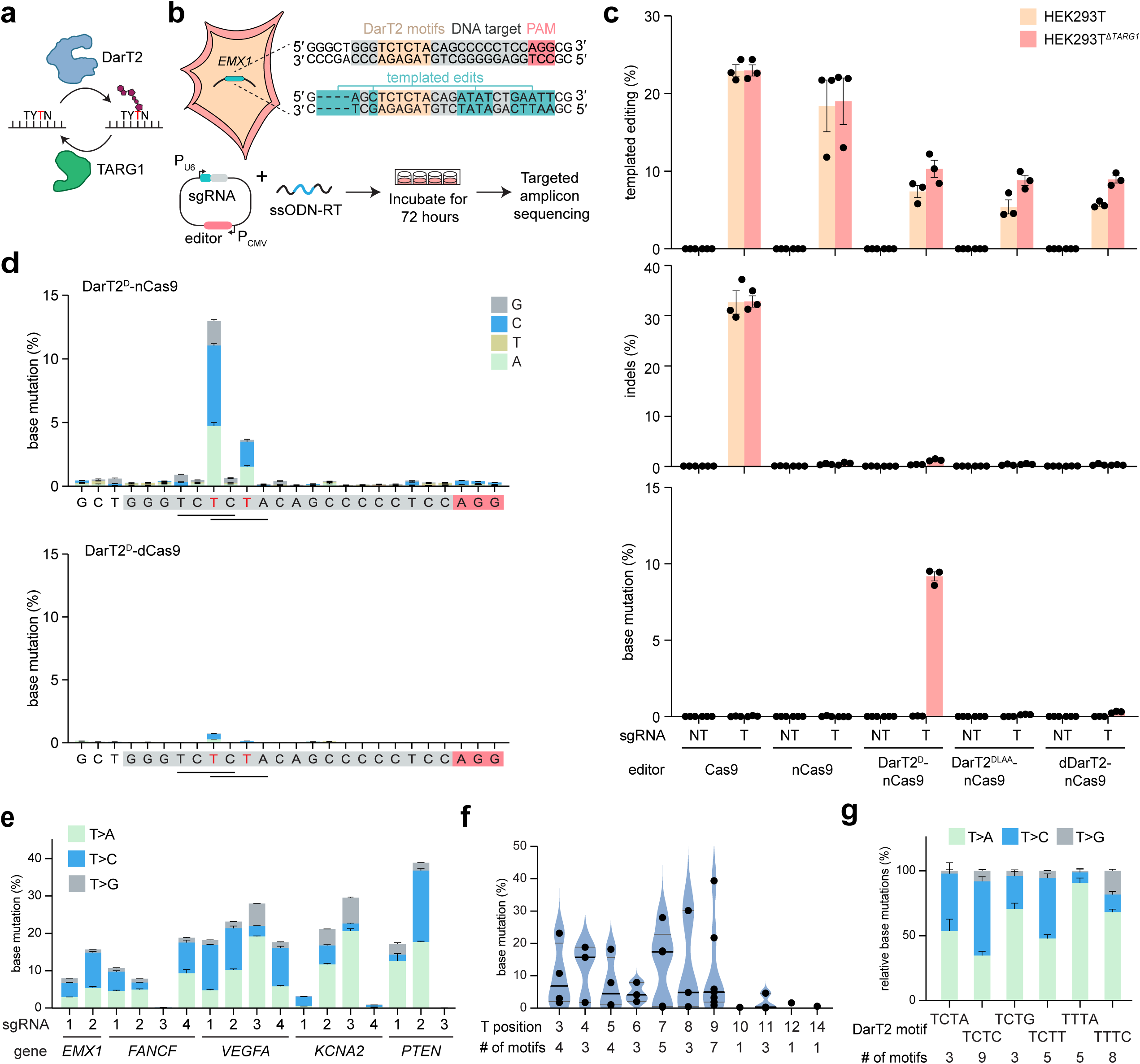
Programmable DNA ADP-ribosylation preferentially drives base mutagenesis in human cells lacking TARG1. **a**, Reversion of ADP-ribosylation of ssDNA in human cells by the TARG1 protein. **b**, Experimental setup for introducing edits in the *EMX1* gene in HEK293T cells using an oligonucleotide repair template (RT). **c,** Extent of templated recombination (top), indel formation (middle) or base mutagenesis (bottom) using *EMX*1 sgRNA1 in HEK293T cells with TARG1 intact (WT) or disrupted (Δ*TARG1*). Bars and error bars represent the mean and SEM of three independent transient transfections without selection or sorting. **d,** Frequency of base substitutions across the sgRNA target in the absence of the oligonucleotide RT. Results are shown with DNA nicking by Cas9 intact (top) or disabled (bottom). **e,** Extent of base mutagenesis of the ADP-ribosylated thymine across 17 target sites in five genes. For d and e, bars and error bars represent the mean and SEM of three independent transient transfections without selection or sorting. **f,** Extent of base mutagenesis based on the relative location of the ADP-ribosylated thymine. Cumulative thymine base editing across 21 sgRNA targets, within 37 5′-TYTN-3′ motifs at positions 3-14 (Position 1 being at the PAM-distal end). Solid black lines represent the median, gray lines represent the quartiles. Each dot represents the mean of three independent transient transfections without selection or sorting for a given sgRNA. **g,** Relationship between the outcome of base mutagenesis and the DarT2 recognition sequence. Distributions were calculated for base mutations occurring at 33 DarT2 recognition motifs across 21 sgRNAs. Bars and error bars represent the mean and SEM of fraction of total values.

Using SpCas9 in HEK293T cells as a baseline, we observed matching extents of templated edits (22%) and indels (32%), with no significant difference in the absence of TARG1 (*p* = 0.99 and 0.94 respectively, n = 3) (**Fig. 4c**). Nicking similarly generated a high level of templated edits whether or not TARG1 was intact (18%), but with minimal indels (0.4%) due to the lack of dsDNA breaks. The append editor with DarT2^D^ also yielded templated edits, with the editing frequency increasing from 7% to 10% by disrupting TARG1. However, no significant differences were observed for append editors with the attenuated DarT2^DLAA^ or with dDarT2 (*p* = 0.32 and 0.33 respectively, n = 3), suggesting that the templated edits were driven primarily through DNA nicking rather than DNA ADP-ribosylation.

At the same time, the ADPr-TA editor with DarT2^D^ yielded 9% base substitutions specifically at the modified thymine within two overlapping DarT2 recognition motifs, but only with TARG1 disrupted (**Fig. 4c**). Base substitutions were negligible with DarT2^DLAA^ (0.2%) or dDarT2 (0.3%), suggesting that higher levels of ADP-ribosylation were necessary to drive editing (**Fig. 4c**). Indel frequencies for ADPr-TAE were slightly elevated over nCas9 with TARG1 disrupted (1.5% vs. 0.9%, *p* = 0.03, n = 3) but still 22-fold lower than that observed with Cas9 (33%) (**Fig. 4c**), indicating that the principal repair outcome of ADP-ribosylation and opposite strand nicking is base mutagenesis. We also observed a low frequency of larger deletions that were elevated with DNA nicking (**Fig. S16**), paralleling observations with BEs^29^. Thus, ADPr-TAE in HEK293T cells drives base mutagenesis similar to that in plants and yeast, but only in the absence of TARG1.

As different oligonucleotide templates revealed reduced templated repair with increased base mutagenesis (**Fig. S17**), we repeated the editing assay without the oligonucleotide template. Base mutagenesis at both modified thymines increased to 16% (**Fig. 4d**), with conversion to either A or C at similar frequencies. Additionally, base mutagenesis was reduced by 20-fold to 0.8% in the absence of DNA nicking, indicating the importance of the nick (**Fig. 4d**). ADP-ribosylation in the absence of opposite-strand nicking would also capture Cas9-independent off-targeting^23^, suggesting that such off-targeting would lead to limited editing despite use of a highly-active DarT2. Probing base mutagenesis beyond this target site, we performed transient transfections without the oligonucleotide template at 16 additional target sites in five genes containing one or more DarT2 recognition motifs (**Figs. 4e and S18**). We observed measurable editing at all but two of these sites, with editing frequencies reaching up to 39% (**Figs. 4e and S18**) and indel frequencies 6-110-fold lower with ADPr-TAE than with Cas9 (**Fig. S19**). Similar trends were observed in U2OS^Δ*TARG*^^1^ cells^28^, with generally lower editing frequencies due to lower transfection efficiencies (**Figs. S20-S21**).

The expanded set of target sites allowed us to explore unique features of base mutagenesis. First, indel frequencies measured by next-generation sequencing or predicted using the Rule Set 2 scoring method^30^ at each target site with Cas9 correlated with base-mutagenesis frequencies (Spearman correlation = 0.80 and 0.58 respectively) (**Fig. S22**). The correlation indicates that efficient targeting and DNA cleavage offer a starting point to identify efficient ADPr-TAE sites. Second, across these sites, editing principally occurred at the modified thymine falling between positions 3 and 9 of sgRNA guide (**Fig. 4f**). For targets with multiple DarT2 recognition motifs, co-occurring mutations were observed 1.1-5.1-fold more frequently than expected if the motifs could be edited independently **(Fig. S23)**. Finally, we noticed distinct mutagenesis distributions that strongly depended on the DarT2 recognition motif (**Fig. 4g**). Specifically, 5′-TCTN-3′ motifs were associated with similar conversion frequencies to A and C. In contrast, 5′-TTTN-3′ were associated with a strong bias toward A, with secondary edits biased toward C (5′-TTTA-3′) or equally split between C and G (5′-TTTC-3′). In total, ADPr-TAE can drive base mutagenesis of thymines in human cells paralleling that observed in yeast and plants, with TARG1 countering the effect of DarT2.

## DISCUSSION

In this work, we explored the impact of appending chemical moieties to target DNA as a distinct yet broad approach for precision editing, what we call append editing. As a first example, we used the bacterial toxin DarT2 to mediate ADP-ribosylation of thymine (abbreviated as ADPr-TAE). When paired with opposite-strand nicking, ADPr-TAE introduced precise edits through homologous recombination in tested bacteria, allowing the creation of templated edits (**Fig. 5**). While this strategy also drove templated recombination in yeast, the predominant outcome was mutagenesis of the ADP-ribosylated thymine. Base mutagenesis was similarly observed in plants and mammalian cells, with a general bias toward substitution to adenine or cytosine (**Fig. 5**). Although the exact underlying repair pathways in eukaryotes remain to be identified (e.g., nucleotide-excision repair, translesion synthesis), homologous recombination can at least be excluded. This divergence in repair pathways contrasts with other genome-editing approaches that engage equivalent repair pathways across organisms and result in similar types of edits, supporting append editing as a distinct entry in the genome editing toolbox.

**Fig. 5:**
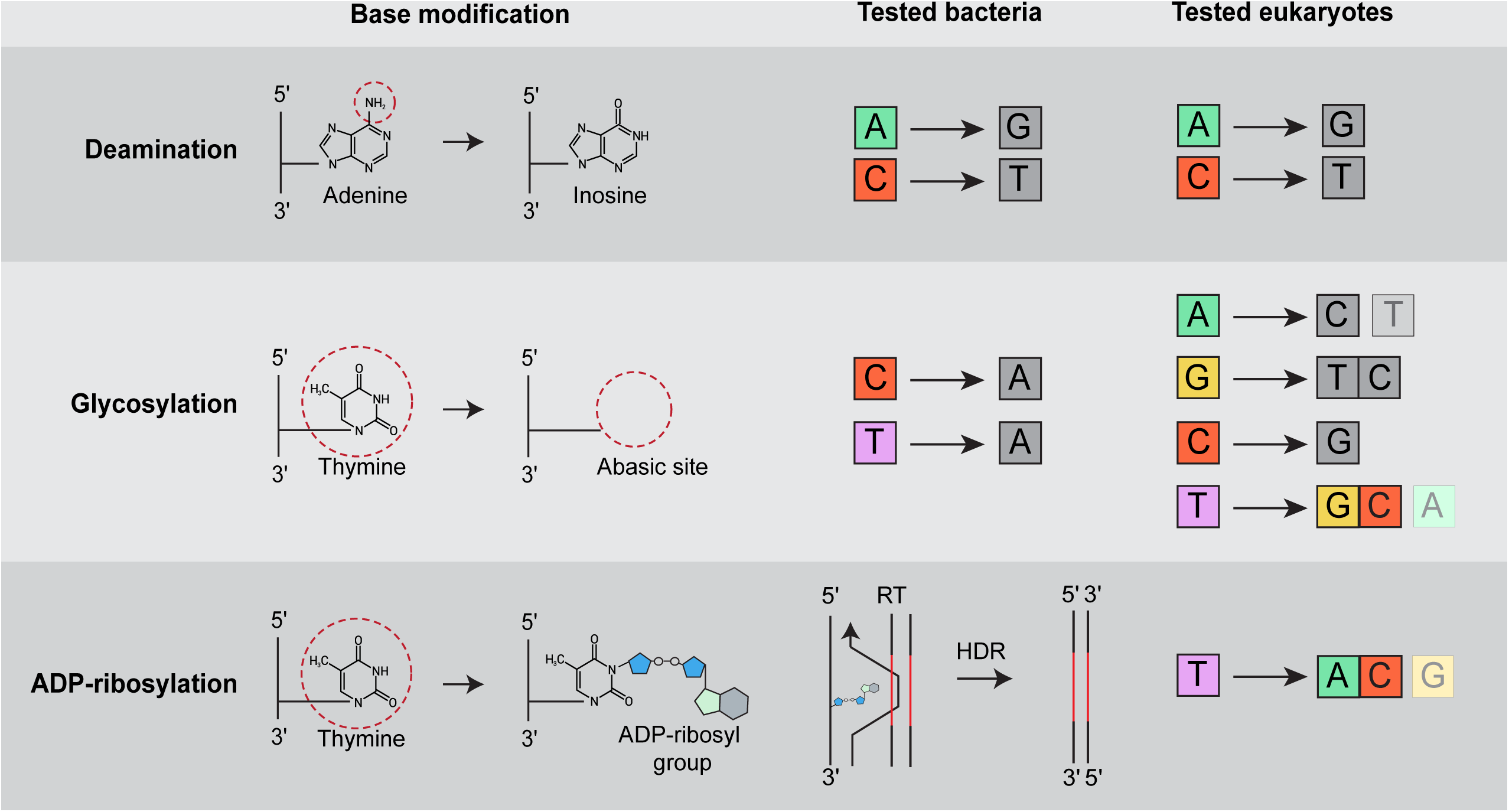
Programmable ADP-ribosylation of thymine generates distinct editing outcomes in bacteria and eukaryotes compared to deaminase and glycosylase base editing. Editors with deaminases include Adenine Base Editors (ABEs)^74^ and Cytosine Base Editors (CBEs)^75^, while editors with glycosylases include A-to-Y Base Editor (AYBE)^36^, Glycosylase Base Editors (GBEs)^76^, Adenine transversion Base Editor (AXBE)^77^, Glycosylase-based Guanine Base Editors (gGBEs/GYBE)^78^, Glycosylase-based Thymine Base Editors (gTBEs), Glycosylase-based Cytosine Base Editors (gCBEs)^40^, Deaminase-Free Thymine Base editor (DAF-TBE), Deaminase-Free Cytosine Base Editor (DAF-CBE)^38^, Thymine DNA glycosylase based editor (TDG), Cytosine DNA glycosylase based editor (CDG)^37^, Thymine base Editor (TBEs)^39^. RT: repair template. HDR: homology-directed repair. Nucleotides representing edits are colored to help compare the glycosylation and ADP-ribosylation of thymine.

ADPr-TAE furthermore offers unique opportunities for genome editing in bacteria (**Fig. 5**) exemplified by the broad range of generated sequence replacements, deletions and insertions. This form of editing did not sacrifice colony counts compared to traditional dsDNA cleavage^31^, offered broader edits without perturbing DNA repair compared to prime editing^32,33^, and omitted fixed scars compared to CRISPR-associated transposons^34^. Given these distinctions, ADPr-TAE is well suited for generating large chromosomal libraries and multiplexed editing or multi-base editing in non-model bacteria^35^.

In yeast, plants, and human cells, ADPr-TAE operates closest to BEs yet offers distinct editing avenues (**Fig. 5**). BEs to date rely on base deaminases or glycosylases that convert T (or A on the opposite strand) into C (adenosine deaminase)^18^, G (adenine glycosylase)^36^ or C/G (thymine glycosylase)^37–40^. In contrast, ADPr-TAE converts T to A or A/C depending on the organism and sequence context. T-to-A editing is particularly unique (**Fig. 5**), where ADPr-TAE could potentially revert 789 of the verified pathogenic SNVs across 355 genes in the ClinVar database^41^ otherwise off-limits through existing thymine base editors. While the current DarT2 recognition motif would capture a fraction of these SNVs (i.e., 30 T-to-A; 447 T-to-C) (**Table S5**), relaxing the motif through ortholog mining or protein engineering could access a greater set. A stringent motif can also be beneficial, such as when reversing pathogenic mutations susceptible to bystander edits. In particular, ADPr-TAE could create a single desired T-to-C edit in a stretch of three thymines (e.g., pathogenic mutation in the third T of 5′-TTTG-3′ (c.103C>T, c.4396C>T, c.4852C>T, c.5188C>T, c.5623C>T, c.742C>T, c.748C>T) or 5′-TTTA-3′ (c.1537C>T, c.3346C>T, c.3673C>T, c.3826C>T, c.4603C>T, c.5473C>T, c.5599C>T)-3′ in the *ATM* gene underlying Ataxia-telangiectasia^42^), while current adenine base editors would generate unwanted edits across the thymines. TARG1 poses an immediate barrier to ADPr-TAE in human cells; however, this barrier could be circumvented with peptide or chemical inhibitors^43^, transient gene silencing such as with RNA interference^44^, or use of dominant-negative inhibitors such as used against mismatch repair^45^.

Beyond ADP-ribosylation of thymine with DarT2, a large number of base-modifying enzymatic domains against any of the four nucleotides could expand append editing. For instance, DarT1 toxins (related to DarT2) and eukaryotic toxins called pierisins (found in cabbage moths) ADP-ribosylate the N2 position of guanine^46,47^, with evidence of base mutagenesis by pierisins in CHO cells^48^. Additionally, bacteria and bacteriophages append unique chemical moieties such as methylcarbamoyl^49^, dPreQ_0_^50^, dADG^51^, glucosyl-5-hydroxymethyl^52^, and 5-hydroxymethyl^53^ to their DNA to block access by anti-phage defenses^54^. The associated enzymatic domains could be further engineered to alter the modified nucleotide, the recognized motif, or the appended moiety as well as enhance editing efficiencies. Interestingly, these examples consistently derive from host-pathogen/parasite interactions that could serve as a plentiful source of such base-modifying domains.

Finally, apart from genome editing, appending chemical moieties to DNA in a targeted manner could facilitate the study of localized versus genome-wide DNA repair. Evaluating the impact of DNA adducts is central to elucidating responsible modes of repair potentially driving mutagenesis and carcinogenesis. To date, introducing such adducts at specific chromosomal sites has proven extremely difficult and laborious^55^. With append editing, specific adducts could be studied in real time^56^ or in conjunction with genome-wide screen of repair pathways^57^ thus uncovering the molecular basis of editing outcomes and probable strategies to shape these outcomes.

## ACKNOWLEDGMENTS

We thank Jimmy Jeske and Rachael Larose for technical support with plasmid cloning and genomic DNA isolation and Ivan Ahel for providing the U2OS^Δ*TARG*^^1^ cell line^28^. The zCas9i MoClo-compatible CRISPR/Cas9 cloning kit with intronized Cas9 was a gift from Sylvestre Marillonnet and Johannes Stuttmann (Addgene kit #1000000171)^58,59^. Genome Analytics Core Unit facility (GMAK) at HZI provided the high-throughput sequencing services. pX330-U6-Chimeric_BB-CBh-hSpCas9 and pSpCas9(BB)-2A-GFP (PX458) were gifts from Feng Zhang (Addgene plasmid # 42230 ^60^; # 48138 ^61^).

## DATA AVAILABILITY

The high-throughput sequencing data have been deposited in the National Center for Biotechnology Information database (BioProject accession PRJNA1149814, https://dataview.ncbi.nlm.nih.gov/object/PRJNA1149814?reviewer=ldg27an527njpmagqvi9use8vj). Source data for all figures are provided in **Table S3** and **Table S4**. There are no restrictions on data availability.

## CODE AVAILABILITY

R scripts used for the analysis of processed NGS data have been deposited on Github (https://gitfront.io/r/Christophe29/9bNjZzbgt6Vk/ADPr-TAE-analysis/).

## AUTHOR CONTRIBUTIONS

**Conceptualization**: S.P.C. and C.L.B. **Methodology**: C.P., D.G., H.V.B., C.K., A.S., J.M.A, C.T., C.L.B. ***In vitro* assays**: C.K. **Bacterial assays**: C.P., H.V.B., I.K., A.M., T.A., A.D.R. **Yeast assays**: C.P., Y.W., T.N., I.S.A. **Plant assays**: K.V., C.Z. **Human cell assays**: D.G., A.K. **Bioinformatic analysis**: D.G, C.T., H.V.B. **Writing manuscript**: C.P., D.G., H.V.B., C.K., C.L.B. **Reviewing and editing manuscript**: all authors, **Figure generation**: C.P., D.G., H.V.B., C.K., C.L.B. **Supervision**: C.P., N.C., E.S., A.S., J.M.A, C.L.B. **Funding acquisition**: A.S., J.M.A., C.L.B.

## METHODS

### Polymerase blocking assays

Wildtype and inactivated (E170A) EPEC DarT2 proteins were expressed using the cell-free myTXTL master mix (Arbor Biosciences). Linear DarT expression templates were amplified from plasmids or ordered as synthetic gene fragments (Integrated DNA Technologies), and contained a T7 promoter and T7 terminator to allow for gene expression (**Table S2**). Cell-free expression was performed in 12 µL reactions, comprising 9 µL of myTXTL master mix, 4 nM of EPEC DarT2 template, 0.4 nM of a T7 RNA polymerase-encoding plasmid, and 4 µM of the RecBCD inhibitor GamS to prevent degradation of the linear DNA templates. The reactions were incubated for 16 h at 29°C.

For ADP-ribosylation of ssDNA templates, the ADP-ribosylation assay was adapted from prior work with slight alterations^13^. Briefly, 5 µL of the TXTL-reaction mix were incubated with 10 µM of the ssDNA oligo, 50 µM NAD^+^, 50 mM Tris-HCl pH 8, 150 mM NaCl, 10 mM EDTA and sterile nuclease free water to reach a final volume of 20 µL, and incubated for 30 min at 30°C. Afterwards, the oligos were separated from the mix using the Oligo clean & concentrator kit (Zymo).

To assess whether DNA ADP-ribosylation blocks DNA polymerases *in vitro*, the DarT-treated oligos were first annealed to the 5′ 6-Fam-tagged primer CKo20 at a final concentration of 10 µM in 1x NEBuffer 2 by heating the mixture to 94°C and gradually cooling it to room temperature. Next, 2 µL of the annealed product were mixed with 0.5 U Klenow Fragment (NEB), 33 µM dNTPs, and 1x NEBuffer 2 in a total volume of 12.5 µL, and incubated for 15 min at 37°C. To stop the reaction, EDTA was added to a final concentration of 10 mM and the samples were incubated at 75°C for 20 min.

To visualize the block of polymerisation, 4 µL of the polymerisation product was mixed with 4 µL loading dye (containing 95% formamide, 0.03% SDS, 18 mM EDTA, 23 µM xylene cyanol, and 19 µM bromophenol blue), and loaded onto a pre-heated denaturing polyacrylamide gel (8 M urea, 20% PAA (19:1)). The gel was run at 250 V for 30 min and visualized under UV light before and after staining with SYBR Gold (Thermo Fisher).

### Microbial strains, handling and growth conditions

All bacterial and yeast strains used in this study are listed in **Table S2**. Unless otherwise specified, *E. coli* TOP10 was used for plasmid cloning and propagation and was grown at 37°C in LB liquid medium (10 g/L tryptone, 5 g/L yeast extract, 10 g/L NaCl) shaking orbitally at 200 rpm, or on LB solid medium (15 g/L agar) at 37°C, containing kanamycin (50 mg/L), carbenicillin (100 mg/L) or chloramphenicol (34 mg/L), when appropriate. The *E. coli kanR*\* strain (CBS-4802) began as strain CB330 (*E. coli* MG1655 P_J23110_-*araFGH* Δ*araBAD*), selected for uniform arabinose induction, to which two chromosomal modifications were made. First, the Δ*lacZ* phenotype (W519*) was generated by CBE-mediated deamination of 5′-ACC-3′ to 5′-ATT-3′ (POS 364749 & 364750 in MG1655), resulting in a premature stop codon; this edit was not used in this work. Second, a defective *kanR* expression construct (*kanR\**) (see **Table S2** for an annotated sequence of the genomic locus) containing a premature stop codon (Q177*) and DarT2 motif 5′-TTTC-3′, was inserted between genes *ybjM* and *grxA* (POS 890463 - 890480 in MG1655) by Red-mediated recombination with Cas9 counterselection^1,31,62^. The resulting *E. coli* MG1655 *kanR*\* strain was used for all assays related to the *kanR*\* gene. The *kanR*\* strain was further used to generate Δ*recA*, Δ*recB*, Δ*recF*, Δ*recT*, Δ*recJ*, Δ*recO*, Δ*xthA*, Δ*mutS* and Δ*uvrA* mutants by Red-mediated recombination^63^. Briefly, transformants of the *E. coli kanR*\* strain carrying pKD46 (encoding λ Red-γ, -β,-exo) were cultured in L-arabinose at 30°C until an OD_600_ of ∼0.6, made electrocompetent as previously described^63^, then transformed with a linear dsDNA template containing 40 nt homology arms to mediate deletion of the target gene. Next, pKD46 was cured from the bacteria by growing them at 37°C, after which the bacteria were made electrocompetent and transformed with pCP20, then grown at 42°C to simultaneously express FLP recombinase and eliminate pCP20. Colonies were then screened for gene deletion by colony PCR and Sanger sequencing. For the substitution assays targeting the *aaaD*, *punR*, *ygcQ* and *yheO* genes, the *E. coli* MG1655 strain was used.

*Salmonella enterica* subsp. enterica serovar Typhimurium str. LT2 was used for all ADPr-TAE assays in *Salmonella* and was regularly grown at 37°C in LB liquid medium shaking orbitally at 200 rpm or on solid LB medium. Carbenicillin (100 mg/L) and chloramphenicol (34 mg/L) were supplemented in the growth medium when necessary.

The *S. cerevisiae* BY4741 (Δ*trp1*, Δ*leu2*) strain was used for all yeast experiments. Unless otherwise specified, *S. cerevisiae* was grown in non-selective liquid YPD medium (20 g/L peptone, 10 g/L yeast extract, and 2% (w/v) D(+)-glucose) or on solid non-selective YPD medium (20 g/L agar). To select for transformants, *S. cerevisiae* cells were grown on solid synthetic defined (SD) medium w/o tryptophan and leucine containing 6.9 g/L yeast nitrogen base without amino acids (Formedium LTD, Cat. # CYN0402), 0.64 g/L complete supplement mixture w/o tryptophan and leucine (Formedium LTD, Cat. # DCS0569), 20 g/L D(+)-galactose (Sigma-Aldrich, Cat. # 15522-250G-R) and 20 g/L agar (Th. Geyer GmbH, Cat. # 214510).

### Plasmid construction

Annotated sequences of all plasmids used in this study are provided in **Table S2**. Unless otherwise specified, general cloning methods such as KLD (KLD Enzyme Mix, Cat. #M0554S) or Gibson assembly (NEBuilder® HiFi DNA Assembly Master Mix, Cat. # E2621X) were used to assemble linear dsDNA fragments into plasmids. Linear dsDNA fragments were amplified with Q5® High-Fidelity 2X Master Mix (NEB, Cat. # M0492L) and purified using the NucleoSpin Gel and PCR Clean-up kit (Macherey-Nagel GmbH, Cat. # 740609.50). Plasmid sequences were verified either by full plasmid sequencing (Plasmidsaurus Inc) or Sanger sequencing (Microsynth Seqlab GmbH).

To generate the append editors expressed in plants, the codon-optimized DNA sequence for DarT2^D^ was commercially synthesized (Twist Bioscience) with a previously reported N7-NLS for expression in *N. benthamiana*^64^, while the zCas9i (*Z. mays* codon-optimized Cas9 coding sequence with 13 introns) was obtained from Addgene (Kit #1000000171)^58^. Both fragments were amplified using the iProof™ High-Fidelity PCR Kit (Bio-Rad, Cat. #1725331). The dDarT, nzCas9i and dzCas9i variants were generated using inverse PCR. Three gRNAs targeting the phytoene desaturase 1 gene (*PDS1*) **(Table S2)** were cloned by annealing complementary oligos into an AtU6 gRNA cassette. Gene fragments were assembled using the GoldenBraid cloning strategy^65^.

### *kanR\** reversion

To assess ADPr-TAE in *E. coli*, an overnight culture of strain CBS-4802 was back-diluted 100-fold, grown to ABS_600_ of 0.6-0.8, then rendered electrocompetent in 10% glycerol. For transformation, 40 μL of electrocompetent cells were mixed with the relevant plasmid(s) and transferred to an ice cold 1-mm electroporation cuvette (Bio-Rad Laboratories, Cat. 1652089). Cells were electroporated using the Gene Pulser Xcell Microbial System (Bio-Rad Laboratories; Cat # 1652662) and the following settings: 1.8 kV, 25 µF, 200 Ω. Next, cells were supplemented with 500 μL of SOC medium and recovered for 1 h at 37°C, shaking orbitally at 200 rpm. Cells were collected by centrifugation at 3000x g, the supernatant was decanted and cells were resuspended in 2 mL induction medium (LB, L-arabinose (2% w/v), carbenicillin (100 mg/L) and chloramphenicol (34 mg/L)) and incubated at 37°C for 16 h, shaking orbitally at 200 rpm. Afterwards, cell cultures were serially diluted in five ten-fold steps in LB, from which 5 μL of each dilution was spotted on LB solid medium containing either carbenicillin and chloramphenicol to select for transformed cells, or carbenicillin, chloramphenicol, and kanamycin to select for transformed and edited cells. The spotted LB solid medium was then incubated for 16 h at 37°C followed by counting colonies.

### Replacement, deletion, and insertion assays in *E. coli*

For the *E. coli* replacement, deletion, and insertion assays at the *kanR*\* locus and the substitution assays at the *aaaD*, *punR*, *ygcQ,* and *yheO* genes, an identical transformation and selection protocol was used as described above. However, after the 16 h incubation in the induction medium, 100 μL of the cell culture was plated on LB solid medium containing carbenicillin and chloramphenicol to obtain single colonies. Single colonies were resuspended in Q5® High-Fidelity 2X Master Mix (NEB, Cat. # M0492L) containing the appropriate primers and subjected to PCR amplification following the instructions of the manufacturer and extending the initial heating step of 98°C to 5 mins to mediate cell lysis and release of genomic DNA. Amplicons were purified and sequenced through Sanger sequencing (Microsynth Seqlab GmbH).

### Growth-based toxicity assay in *E. coli*

The growth-based toxicity assay began by rendering strain CBS-5301 electrocompetent. Next, 9 fmol of plasmid CBS-4808 was transformed into strain CBS-5301 using the electroporation conditions described above. Transformants were recovered in 500 µL of SOC medium for 1 h at 37°C, shaking orbitally at 200 rpm, then plated on LB solid medium supplemented with carbenicillin and incubated for 16 h at 37°C. Next, a single colony was inoculated into 2 mL LB medium containing carbenicillin, grown until an OD_600_ of 0.6, then made electrocompetent following the protocols described above. A second round of transformation was performed, using one of nine different editor plasmids (CBS-6738/-6739/-6741/-6742/-6743/-6744/-6745/4781/-4800), following the electroporation protocol described above. Transformed cells were allowed to recover in 500 µL SOC medium for 1 h at 37°C shaking orbitally at 200 rpm, plated on LB solid medium supplemented with carbenicillin, chloramphenicol and glucose (20 mM), and incubated for 16 h at 37°C. Three individual colonies from each of the nine resulting strains (**Table S2**) were then used to inoculate a 96 deep-well plate (Greiner Bio-One Cat. # 780271), containing 400 µL of LB medium supplemented with carbenicillin, chloramphenicol and glucose (20 mM) and covered with an adhesive gas-permeable membrane (Thermo Scientific, Cat. # 241205). After incubating the deep-well plate for 16 h at 37°C, the cell cultures were adjusted to an OD_600_ equal to 0.1 using LB supplemented with carbenicillin, chloramphenicol, and L-arabinose (0.2% w/v) in a new 96-well plate, reaching a final volume of 200 µL. The 96-well plate was then measured every 3 minutes over 12 h at 37°C for absorbance at 600 nm on a BioTek Synergy Neo2 plate reader, shaking at 500 rpm.

### Non-selective editing at *kanR*\*

Transformations were performed as described above, however after the 16 h incubation in induction medium, the cultures were centrifuged, the medium was discarded, and genomic DNA was isolated using Wizard Genomic DNA Purification Kit (Promega, Cat. # A1120). The kanR site was then amplified through PCR using the primer pair HBo-314 and HBo-315 and the Q5® High-Fidelity 2X Master Mix (NEB, Cat. # M0492L) for 25 cycles. Resulting amplicons were sequenced with Nanopore sequencing (Eurofins Genomics Germany GmbH). For data analysis, FASTQ sequencing data files were aligned to a FASTA file of the unedited amplicon using MiniMap2 with option “map-ont”^66^. Samtools was used to convert the sequence alignment/map (SAM) files into binary alignment/map (BAM) files, while concurrently sorting and indexing^67^. All further analysis was performed using R, after calling libraries tidyverse and GenomicAlignments^68^. A function was defined to take BAM files as an argument, then extract all alleles aligned to the 8 nucleotide region of the templated edit as a list of characters. This function was applied to all BAM files to generate lists of alleles, which were tallied and compiled into a single data frame in long table format. Next, alleles were defined as unedited, edited, or ambiguous, and the fraction of each observation was computed. Samples were then grouped by editor and repair plasmids, after which the mean and standard deviation were computed, then used to generate the bar plot. Further analysis was undertaken to search for base mutations at the ADPr site. The list of alleles in the initial data frame was filtered to retain only records containing a T-to-V mutation at the ADPr target position, but otherwise match the reference allele. Records were grouped by sample, SNVs were tallied, after which each was divided by the total number of observed alleles and multiplied by 100, to obtain the percent of base mutations amongst all sequencing reads.

### Whole genome off-target assay in *E. coli*

For identifying whole genome off-target mutations, strain CBS-4802 was grown from a single colony in LB medium and made electrocompetent as described above. Electrocompetent CBS-4802 was then co-transformed with equimolar amounts (9 fmol) of CBS-6746 and one of several editor plasmids (CBS-3130/-6738/-6740). Transformants were recovered in 500 μL SOC for 1 h at 37°C shaking orbitally at 200 rpm, after which the growth medium was replaced with 2 mL of LB, supplemented with carbenicillin, chloramphenicol, and L-arabinose (0.2%), followed by incubation at 37°C for 16 h shaking orbitally at 200 rpm. Next, the cultures were streaked onto LB solid medium supplemented with carbenicillin and chloramphenicol and incubated for 16 h at 37°C in order to obtain individual colonies. Three colonies from each condition were in 2 mL of LB medium supplemented with carbenicillin and chloramphenicol, and cultured for 16 h at 37°C.

After incubation, cultures were centrifuged and the cell pellets were subjected to genomic DNA isolation using the Wizard Genomic DNA Purification Kit (Promega, Cat. # A1120). Isolated genomic DNA was fully sequenced using Nanopore sequencing (Plasmidsaurus Inc). For data analysis, FASTQ sequencing data files were aligned to a FASTA file of *E. coli* MG1655 (GenBank: U00096.3), using Minimap2, with the “map-ont” option^66^. Samtools was used to convert the sequence alignment/map (SAM) files into binary alignment/map (BAM) files, while concurrently sorting and indexing^67^. Clair3 was run on the GalaxyEU server, to call variants^69,70^. Bcftools was used to query the variant call format (VCF) files for POS, REF, ALT, DP, and AF fields, and export the results into a comma-separated values (CSV) file^71^. The sequencing depth at all positions in all BAM files was calculated by Samtools, and exported as a CSV file. All further analysis was performed in R after loading library tidyverse^68^. CSV files were loaded into a long format dataframe. This dataframe was then filtered with the following steps. 1) SNVs were retained, by filtering for records that contain only a single character in the REF and ALT fields. 2) SNVs already present in the parent strain were eliminated, by filtering for records containing POS field values not found in parent strain POS field values. 3) SNVs mapped to regions known to have been modified during the creation of strain CBS-4802 were eliminated, by filtering for records with POS field values not present in said regions. 4) Records were filtered for AF field values greater than or equal to 0.25. 5) SNVs observed at a sequencing depth greater than or equal to the lowest quartile of all BAM files (Q1>=34) were retained. 6) All SNVs were re-coded to C>D and T>V, tallied, then used to generate a heatmap.

### Editing assays in *S. enterica*

Electrocompetent *S. enterica* cells were transformed with 9 fmol of plasmid CBS-4800 and recovered in 500 μL SOC medium following an identical protocol as the one described above for *E. coli*. After recovery, the cells were collected through centrifugation at 3,000x g, the supernatant was decanted, and the cell pellet was resuspended in 100 μL of LB medium. The cell suspension was plated on LB solid medium containing chloramphenicol (34 mg/L) and incubated at 37°C for 16 h. After incubation, a single colony was selected and used to create electrocompetent *S. enterica* cells harboring plasmid CBS-4800, following the protocol described above. Then, 22 fmol of the plasmids containing the repair template and the targeting (T) sgRNA (**Table S2**) were transformed in triplicate through electroporation into *S. enterica* cells harboring plasmid CBS-4800. The cells were recovered in 500 μL SOC medium, collected through centrifugation at 3,000x g, the supernatant was decanted, and the cell pellet was resuspended in 2 mL of induction medium (LB, 2% (w/v) L-arabinose, 100 mg/L carbenicillin and 34 mg/L chloramphenicol) and grown at 37°C for 16 h, shaking orbitally at 200 rpm. 100 μL of the cell culture was plated on LB solid medium containing carbenicillin and chloramphenicol to obtain single colonies. Colonies were resuspended in Q5® High-Fidelity 2X Master Mix (NEB, Cat. # M0492L) containing the appropriate primers and subjected to PCR amplification following the instructions of the manufacturer and adding an initial heating step of 98°C for 5 min to mediate cell lysis and release of genomic DNA. Amplicons were then purified using the NucleoSpin Gel and PCR Clean-up kit (Macherey-Nagel GmbH, Cat. # 740609.50) and sequenced through Sanger sequencing (Microsynth Seqlab GmbH).

### Templated editing assays in *S. cerevisiae*

*S. cerevisiae* BY4741 (Δ*trp1*, Δ*leu2*) cells were co-transformed with two plasmids, one bearing either of the editor variants (DarT^DLAA^-nCas9 or dDarT-nCas9) and the other bearing a 6 bp substitution template flanked by 294-bp (upstream) and 232-bp (downstream) homology arms along with either an *FCY1* targeting (T) sgRNA or a non-targeting (NT) sgRNA (**Table S2**), following the lithium acetate method as previously described^72^.

Briefly, single *S. cerevisiae* colonies were inoculated into 2 mL liquid YPD medium (20 g/L peptone, 10 g/L yeast extract, 2% (w/v) D(+)-glucose) and grown for 16 h at 30°C, shaking at 200 rpm on a rotary shaker. The cells were diluted to an OD_600_ equal to 0.5 in 50 mL of YPD medium and cultured again at 30°C, shaking at 200 rpm, until the cells reached an OD_600_ equal to 2. The cells were then harvested by centrifugation at 3,000x g for 5 min, the supernatant was decanted and the pellet was resuspended in 25 mL of sterile water. The centrifugation and resuspension step was repeated followed by another centrifugation at 3,000x g for 5 min and resuspension in 1 mL of sterile water. The cell suspension was then centrifuged for 30 s at 13,000x g, the supernatant was discarded and the pellet was resuspended in 1 mL of sterile water. 100-μL aliquots were distributed in 1.5 mL sterile Eppendorf tubes, and the cells were collected by centrifugation at 13,000x g for 30 s. The supernatant was decanted and the cell pellet was resuspended with 336 μL of transformation mix (240 μL of PEG 3350, 36 μL of 1 M LiAc, 50 μL of 2 mg/mL single-stranded carrier DNA), plasmid DNA (500 ng of each plasmid) and sterile water to reach a final volume of 360 μL. The suspension was incubated at 42°C for 40 min, after which it was centrifuged at 13,000x g for 30 s. The supernatant was decanted, the cell pellet was resuspended in 1 mL of YPD and the cell suspension was incubated for 3 h at 30°C. Cells were collected by centrifugation at 13,000x g for 30 s and washed twice with 1 mL of SD medium to remove any residual YPD medium. Finally, the cell pellet was resuspended with 100 μL of SD medium, plated on solid SD medium without tryptophan and leucine and containing D-galactose, and incubated at 30°C for 3 days or until colonies were visible.

Resulting colonies were collected with a sterile 10 μL pipette tip and resuspended in 10 μL sterile 0.02 M NaOH, boiled at 99°C for 10 min and centrifuged for 10 s at maximum speed in a microcentrifuge. 1 μL of the supernatant was used as template for PCR using the Q5® High-Fidelity 2X Master Mix (NEB, Cat. # M0492L) and the primer pair prCP222-prCP223 to amplify *FCY1* (**Table S2**). The resulting PCR product was purified using the NucleoSpin Gel and PCR Clean-up kit (Macherey-Nagel GmbH, Cat. # 740609.50), following the manufacturer’s instructions. The final product was sequenced through Sanger sequencing (Microsynth Seqlab GmbH). Sequence alignment was performed using the online MAFFT algorithm^73^.

### Base mutation assays in *S. cerevisiae*

*S. cerevisiae* BY4741 (Δ*trp1*, Δ*leu2*) cells were co-transformed with two plasmids, one bearing either of the editor variants (DarT^DLAA^-nCas9 or dDarT-nCas9) and the other bearing either of the targeting (T) sgRNAs for *FCY1*, *ALP1* or *JSN1*, or a non-targeting (NT) sgRNA (**Table S2**), following identical procedures as described above. Resulting colonies were screened through colony PCR as described above, and the primer pairs prCP222-prCP223, prCP445-prCP446 and prCP441-prCP442 were used to amplify *FCY1*, *ALP1* and *JSN1*, respectively (**Table S2**). The resulting PCR products were sequenced through Sanger sequencing, and sequence alignment was performed using the MAFFT algorithm^73^.

### Base mutation assays in *N. benthamiana*

*N. benthamiana* seeds were germinated in soil and transplanted at one-week-old stage to 24 cell nursery flats, one plant per cell, and grown at 23°C under a 16-h-light and 8-h-dark cycle in Sungro horticulture professional grow mix mixed 1:1 with Jolly gardener Pro-line C/B growing mix (Sungro).

Plasmids were used to electroporate *Agrobacterium tumefaciens* strain GV3101 using Bio-Rad GenePulser electroporator with the following conditions: 1.8 kV, 100 Ω, and 25 µFD. Single colonies were inoculated in LB medium containing spectinomycin (100 µg/mL), rifampicin (50 µg/mL), and gentamicin (50 µg/mL) for 16 h at 28°C with orbital shaking at 200 rpm. Cultures were then centrifuged and resuspended in infiltration medium (10 mM MgCl_2_ and 100 µM Acetosyringone) to reach an OD_600_ of ∼0.1. Following, the resuspended cultures were combined in a 1:1 ratio with an *A. tumefaciens* strain containing *p19* (a suppressor of gene silencing) and were infiltrated into the leaves of four-week-old plants using a 1-mL needleless syringe. The infiltrated plants were then recovered overnight in the dark and grown for 7 days using conditions mentioned above.

### Next-Generation Sequencing in *N. benthamiana*

Leaf tissues were harvested 7 days post-infiltration using a standard hole-punch and collected in 1.5 mL tubes containing ∼100 µL of 1 mm glass beads. Disks from four leaves (one disk per leaf) were pooled to create each biological replicate. The samples were frozen at −80°C for 24 h, after which the tissue was ground using a Vivadent shaker for 5 s followed by resuspension in CTAB buffer (1.4 M NaCl, 20 mM EDTA, pH 8, 100 mM Tris-HCl, pH 8, 3% CTAB (cetyltrimethylammonium bromide)). Cellular DNA was then extracted using chloroform and isopropyl alcohol followed by a 70% ethanol wash.

The targeted region was amplified with optimized primers and PCR conditions, using iProof™ High-Fidelity PCR Kit (Bio-Rad, Cat. #1725331). The products were purified using 4 µL of ExoSAP-IT™ PCR Product Cleanup Reagent (Applied Biosystems, Cat.# A55242) at 37°C for 15 min followed by inactivation at 80°C for 15 min. A second amplification was performed with iProof polymerases to introduce unique Illumina Barcodes and libraries were purified using QIAquick Gel Extraction Kit (Qiagen).

The concentration for each library was measured using Qubit fluorometer (Invitrogen) and equimolar amounts were pooled alongwith 120 pM phiX control library corresponding to 8% of the final volume. 20 μL of the pooled library was loaded into the iSeq 100 (Illumina) and the run was performed in accordance with iSeq 100 Sequencing System Guide. Sequencing data analysis was performed as mentioned for mammalian cells.

### Mammalian cell cultures and transfection

HEK293T cells were purchased from ATCC (CRL 11268) and U2OS^Δ*TARG*^^1^ cell lines were a gift from the Ahel lab. Unless otherwise mentioned, all cell lines were maintained using Dulbecco’s modified Eagle’s medium (Life Technologies) supplemented with 10% (v/v) fetal bovine serum (Corning and BANF Biotrend), 1x penicillin streptomycin (Life Technologies) and 2 mM L-glutamine. The cultures were incubated in humidified incubators at 37°C with 5% CO_2_.

For generating HEK293T^Δ*TARG*^^1^ cell line, cells were transfected with plasmids containing WT-SpCas9 and *TARG*1 sgRNA^28^ (**Table S2**) using Lipofectamine 3000™ (Invitrogen, Cat.# L3000008) according to the manufacturer’s instructions. 48 h post-transfection, cells were diluted and seeded in 96-well plates at a density of 3 cells/well. Colonies were observed after 7 days and wells with single colonies were selected. Selected clones were tested for *TARG1* site disruption through Sanger sequencing followed by Western Blotting **(Fig. S15.)** with anti-*TARG1* antibody (Fisher Scientific, Cat.# 25249-1-AP)^28^ and anti-beta-actin antibody (Life technologies, Cat.# MA5-15739-HRP) as the housekeeping control.

For templated editing assays in HEK293T (WT and Δ*TARG1*) cell line, 65,000 cells/well were seeded onto tissue culture treated 24-well plates (Corning) and incubated at 37°C with 5% CO_2_ under humidified conditions. 24 h later, 50 fmol of each plasmid was co-transfected with 750 fmol of ssODN repair templates using 1.12 μL of lipofectamine 3000™ reagent and 1 μL of P3000. For base mutagenesis assays, 500 ng of each plasmid was transfected, following the same conditions as mentioned above. The medium was refreshed 24 h post transfection and the cultures were harvested 72 h post transfection.

For base mutagenesis assays in the U2OS^Δ*TARG*^^1^ cell line, 1.3 x 10^5^ cells were seeded and 1 μg of plasmid DNA, 1.5 μL of lipofectamine 3000™ reagent, and 2 μL of P3000 were used for transfection. Media change and sample harvest were performed similar to HEK293T cells.

### Next-generation sequencing for mammalian cells

Genomic DNA was isolated from harvested cells using PureLink™ Genomic DNA Mini Kit (Life Technologies, Cat. # K182002). Specific primers were used to amplify the targeted region using Q5® High-Fidelity 2X Master Mix (NEB, Cat. # M0492L) through 27 cycles. The PCR product was purified using NucleoSpin Gel and PCR Clean-up Kit (Macherey-Nagel, Cat. #740609) and was used as a template in KAPA HiFi HotStart ReadyMix (Roche Diagnostics, Cat. # KK2602) to introduce Illumina adapter sequences within 15 PCR cycles. The KAPA-PCR products were cleaned using Agencourt® AMPure® XP magnetic beads (Beckman Coulter, Cat. # A63881) and 200 ng of this product was used as template for a second PCR with KAPA Ready Mix to introduce Illumina Barcodes through 10 PCR cycles followed by cleanup using magnetic beads as mentioned before. PCR products were screened at each step for correct fragment length using agarose gel electrophoresis. The libraries were pooled in equimolar amounts and at least one million reads were generated for each sample using NovaSeq™ 6000 and the demultiplexed data was analyzed by CRISPResso2. Default parameters were used to perform the analysis except when quantifying indel and HDR frequencies for templated editing, in which case a plot_window_size = 30 was used. Allelle_frequency_table_around_sgRNA.txt files generated by CRISPResso2 were used within R-scripts (https://gitfront.io/r/Christophe29/9bNjZzbgt6Vk/ADPr-TAE-analysis/) to further quantify Base mutation frequencies as the total percentage of reads containing a nucleotide different from the reference read.

### Statistical analyses

For assays involving *kanR* reversion on solid medium (**Fig. 1g, 2b)**, unpaired, two-tailed, Welch’s t-tests were performed on log-normal data. Figure error bars display standard deviation. For the non-selective editing experiment (**Fig. S3**), a one-way ANOVA was performed to test for the effect of editor-sgRNA combinations on the percent of reads showing a SNV at the target thymidine. For the assay involving deletion strains in *E. coli* (**Fig. 2e**), unpaired, one-tailed, Welch’s t-tests were performed on log-normal data. Figure error bars display standard deviation. For short-read NGS data (**Fig. 3f, 3g, 4c, 4d, 4e, 4g**), unpaired, two-tailed, Welch’s t-tests were performed. Figure error bars display standard error of the mean. For the editing window experiment **(Fig. 4f)** median and quartiles of each group are displayed. Related p-value calculations can be found in **Table S3** and **Table S4**.

