## Supplementary Information for "Targeted DNA ADP-ribosylation triggers templated repair in bacteria and base mutagenesis in eukaryotes"

### SUPPLEMENTARY FIGURES

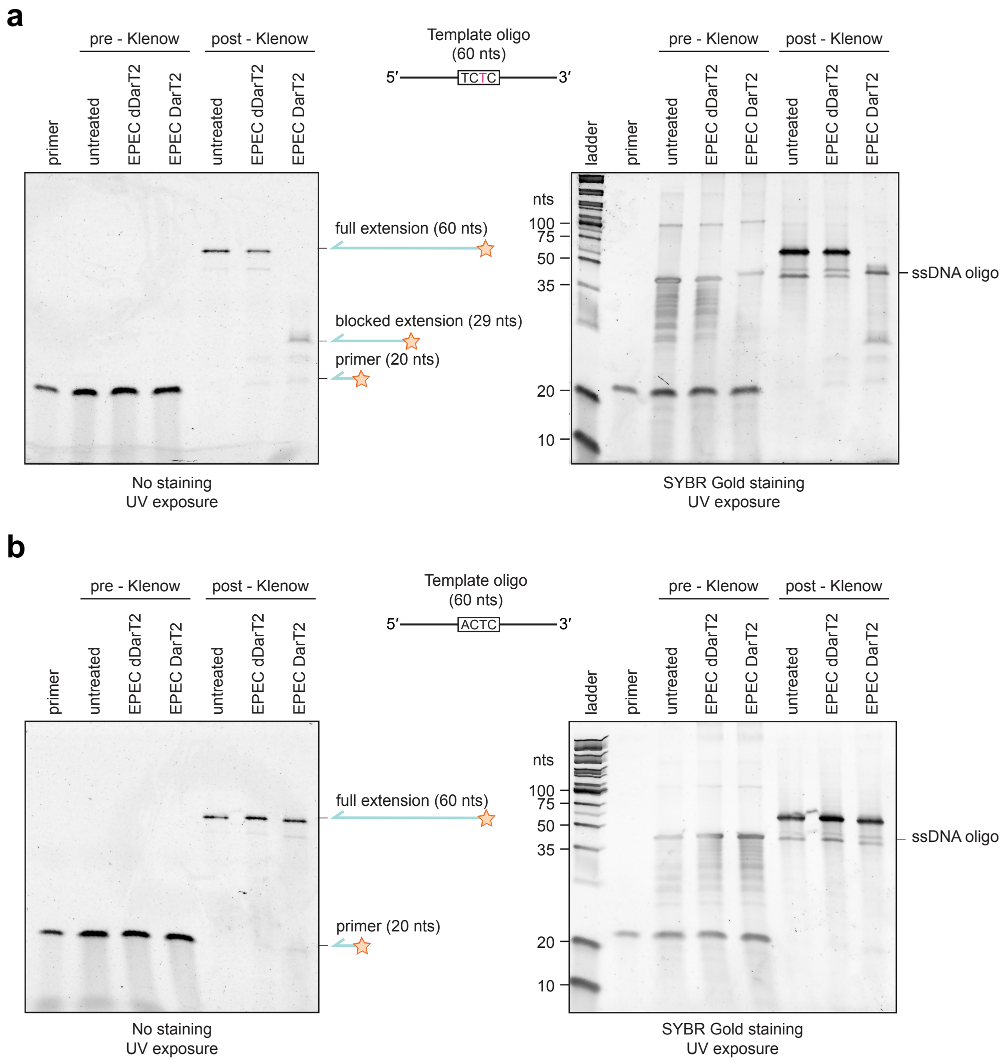

**Fig. S1. ADP-ribosylation of a 5′-TCTC-3′ motif on ssDNA by cell-free expressed EPEC DarT2 blocks DNA polymerisation by *E. coli* Klenow fragment.** ssDNA oligos harboring a 5′-TCTC-3′ (**a**) or 5′-ACTC-3′ (**b**) motif were treated with wildtype or attenuated (dDarT) EPEC DarT2, annealed with a 3′-complementary Fam-tagged oligo, and treated with *E. coli* DNA Polymerase I (Klenow Fragment). Primer extension products were run on denaturing polyacrylamide gels and imaged under UV light before (left, only showing elongated Fam-tagged primer) and after (right, imaging all DNA) SYBR Gold staining. Images are representative of triplicate independent experiments. dDarT2: EPEC DarT2 with the inactivating E170A mutation.

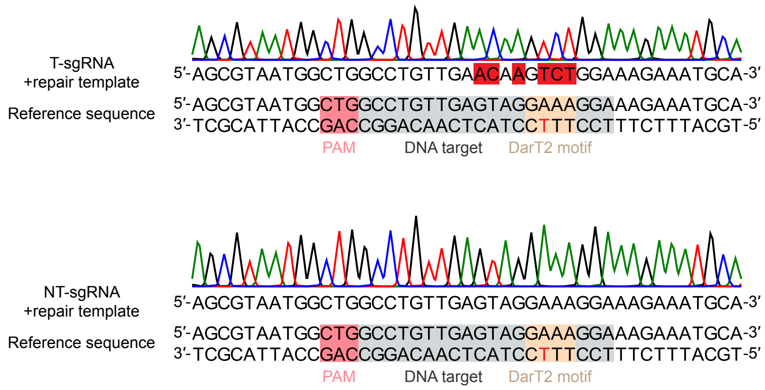

**Fig. S2. Representative Sanger sequencing results after targeting the *kanR** site with DarT2$^{D}$-ScnCas9 in *E. coli* in the presence of a repair template.** The target thymine (T) in the DarT2 motif is labelled in red. The mutated bases are highlighted in red. Results are shown for a screened kanamycin-resistant colony under targeting conditions (top) and a colony under non-targeting conditions (bottom).

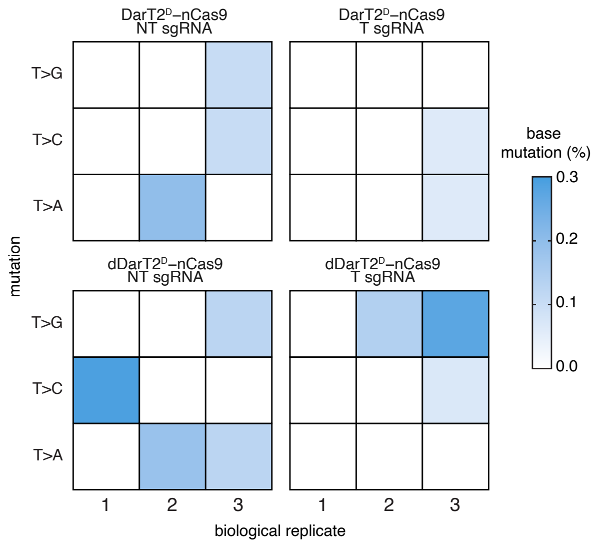

**Fig. S3. Assessing single-nucleotide variants at the ADP-ribosylated thymine targeted with the specified ScCas9 ADPr-TA editor.** Aligned reads are quantified that match the unedited sequence at all positions except at the ADP-ribosylated position as a percent of all aligned reads. Each biological replicate starts from a separate transformation.

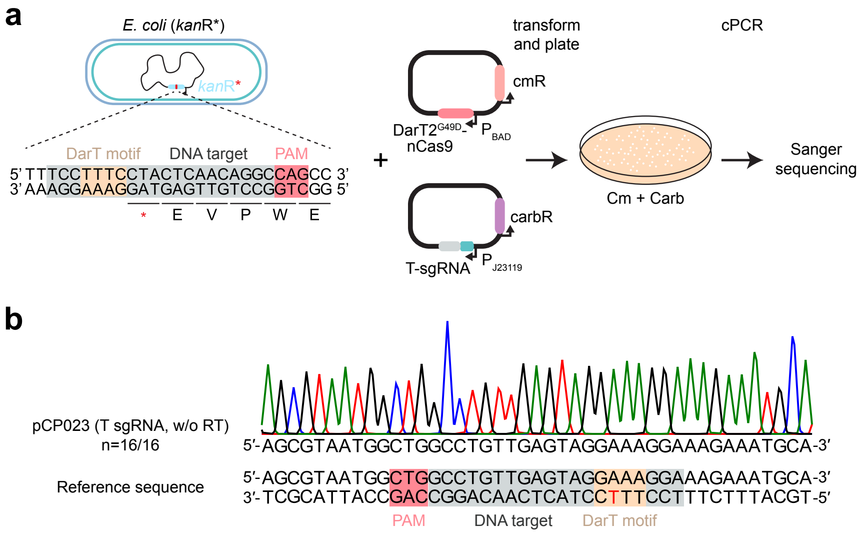

**Fig. S4. Representative genotype after targeting the *kanR** site with DarT2$^{D}$-ScnCas9 in the absence of a repair template. a.** Experimental setup for targeting the kanamycin resistance gene (*kanR**) in *E. coli* using the DarT2$^{G49D}$-nCas9 editor and a targeting (T) sgRNA, without a repair template. **b**. Representative Sanger sequencing chromatogram of 16 sequenced colonies.

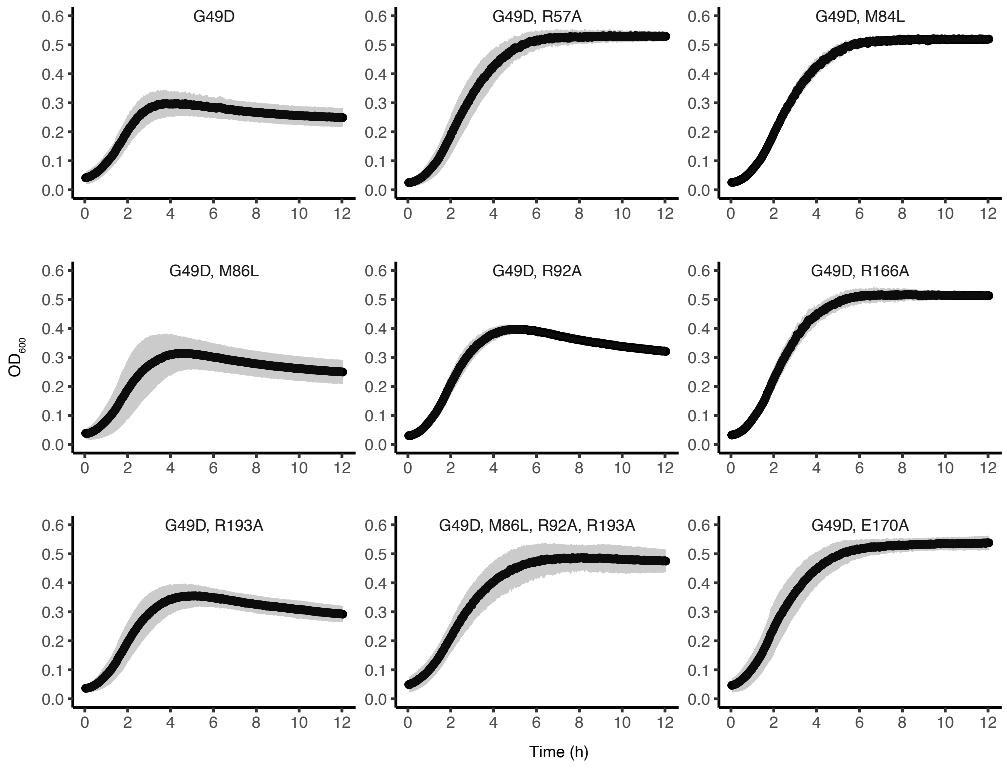

**Fig. S5. Growth curves of *E. coli* Δ*recA* strains transformed with different attenuated variants of DarT2 fused to ScnCas9.** Optical density measurements of cultures from a plate reader over the course of 12 hours, with absorbance measured at 600 nm and absorbance of the culture medium subtracted. Plots show the mean of triplicates in black, and the standard deviation of triplicates in grey.

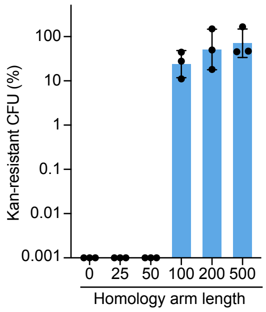

**Fig. S6. Impact of homology arm length on reverting *kanR** with DarT2$^{DLAA}$-ScnCas9 in *E. coli*.** The indicated lengths (in bp) were the same upstream and downstream of the 8-bp edit (e.g., 25 bp means 25-bp homology upstream and 25-bp homology downstream of the edits). Bars and error bars represent the mean and s.d. of three independent experiments started from separate transformations.

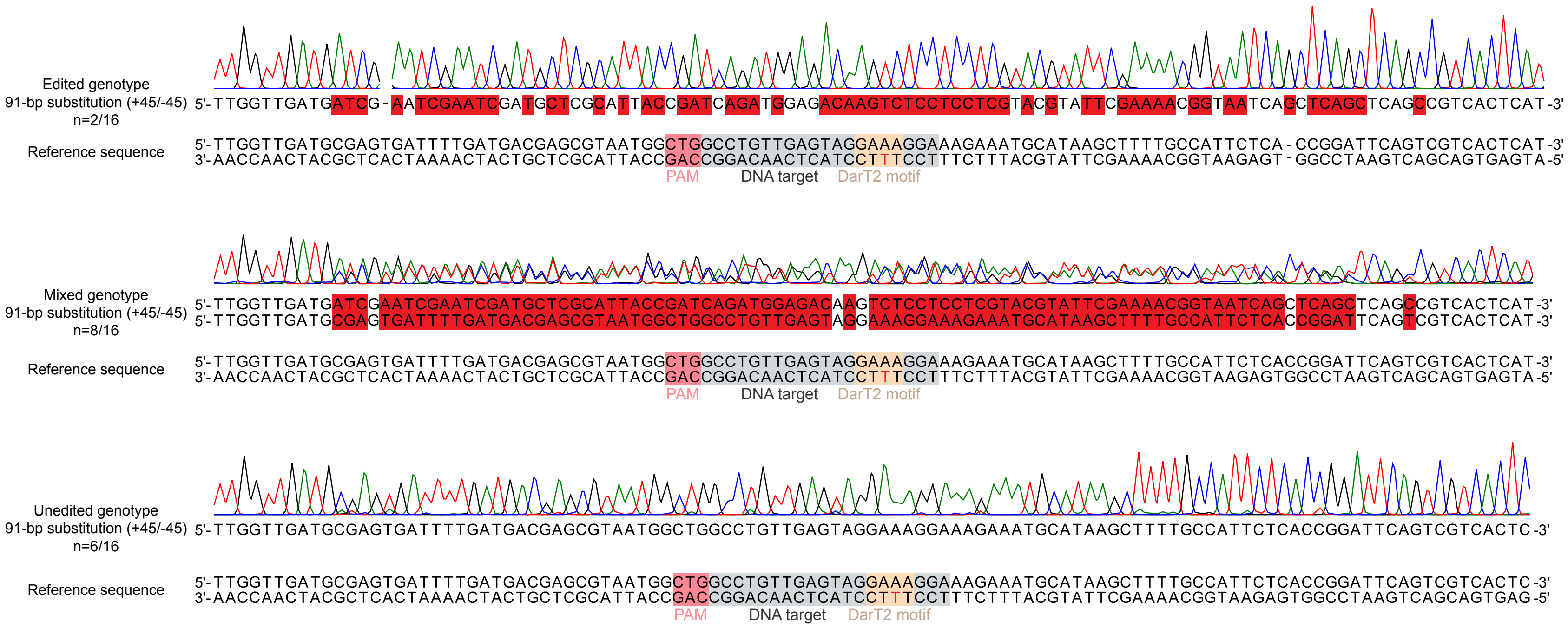

**Fig. S7. Examples of edited, mixed, and unedited genotypes for sequence replacement with DarT2DLAA-ScnCas9 targeting *kanR** in *E. coli*.** Representative Sanger sequencing results from *E. coli* colonies transformed with a plasmid that contained a repair template to mediate the replacement of 91 bp at the *kanR** site; 45 bp upstream and 45 bp downstream (+45/-45) of the ADP-ribosylated thymine.

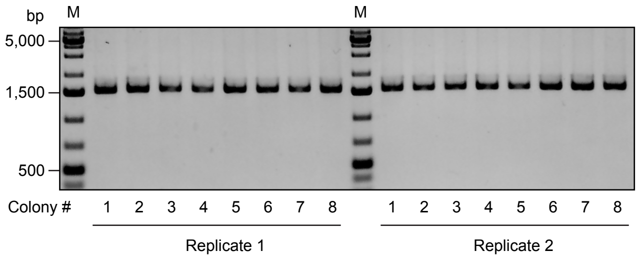

**Fig. S8. Colony PCR screening for 500-bp insertion at the *kanR** target site in *E. coli* following targeting with DarT2$^{DLAA}$-ScnCas9 in the presence of the repair template.** Two replicate transformations were used. Eight colonies from each replicate were screened through colony PCR using primers HBo312 and HBo313. M, marker. Results relate to the 500-bp insertion in Figure 2h.

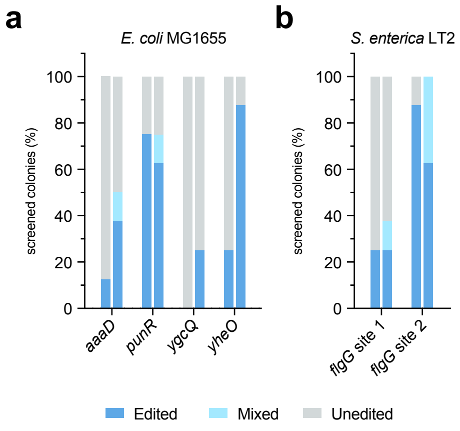

**Fig. S9. ADPr-TAE with DarT2$^{DLAA}$-ScnCas9 at different sites in *E. coli* MG1655 and at two sites in the *flgG* gene in *S. enterica* LT2. a.** ADPr-TA editing through substitution at the *aaaD*, *punR*, *ygcQ,* and *yheO* genes of *E. coli* MG1655. All substitutions introduced silent mutations. **b.** ADPr-TA editing through substitution (*flgG* site 1) or deletion (*flgG* site 2) at the *flgG* gene of *S. enterica* LT2. All experiments for *E. coli* and *S. enterica* were performed in duplicate, and eight colonies were screened through colony PCR and Sanger sequencing per duplicate experiment.

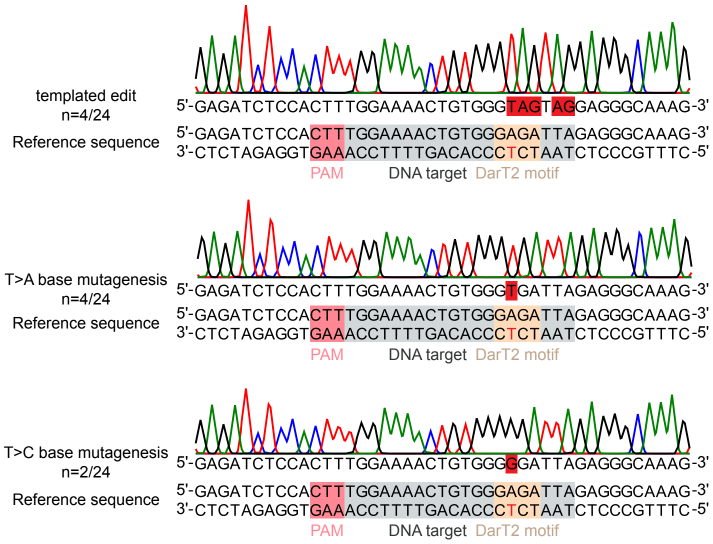

**Fig. S10. Representative genotypes after targeting the *FCY1* gene with DarT2$^{DLAA}$-ScnCas9 in *S. cerevisiae* in the presence of a repair template.** Top, representative Sanger sequencing chromatogram showing the templated edit. 24 colonies were screened, out of which four contained the indicated genotype. Middle, representative Sanger sequencing chromatogram showing T-to-A mutagenesis of the target T (labelled red) in the 5′-TYTN-3′ DarT2 motif. Twenty-four colonies were screened, out of which four contained the T-to-A mutation. Bottom, representative Sanger sequencing chromatogram showing a T-to-C mutagenesis of the target T (labelled red) in the 5′-TYTN-3′ DarT motif. 24 colonies were screened, out of which two contained the T-to-C mutation. The mutated bases are highlighted in red.

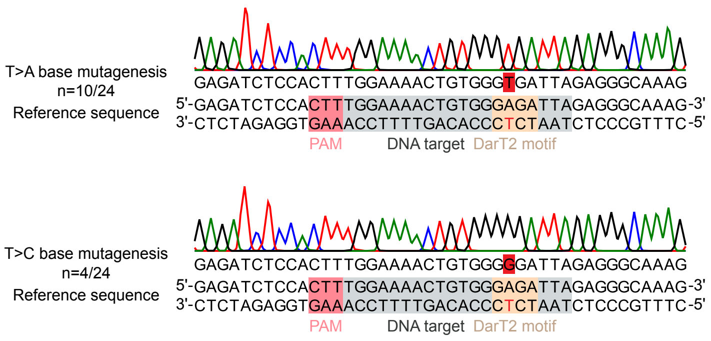

**Fig. S11. Representative genotypes after targeting the *FCY1* gene with DarT$^{DLAA}$-ScnCas9 in *S. cerevisiae*, in the absence of a repair template.** See Fig. S10 for details.

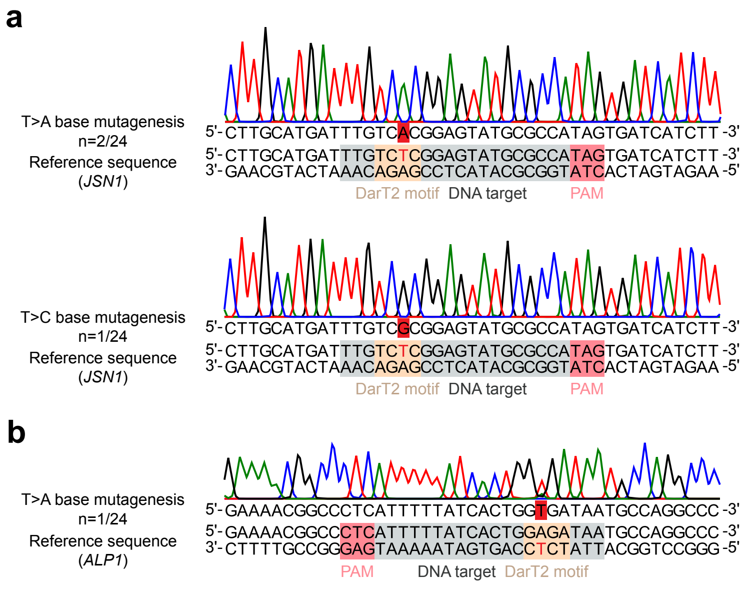

**Fig. S12. Representative genotypes after targeting the *JSN1 or ALP1* genes in *S. cerevisiae* with DarT$^{DLAA}$-nCas9 in the absence of a repair template. a,** Top: representative Sanger sequencing result showing T-to-A mutagenesis of the target T (labelled red) in the 5′-TYTN-3′ DarT2 motif. 24 colonies were screened, out of which two contained the T-to-A mutation. Bottom: representative Sanger sequencing result showing T-to-C mutagenesis of the target T (labelled red) in the 5′-TYTN-3′ DarT2 motif. 24 colonies were screened, out of which one contained the T-to-C mutation. **b,** Representative Sanger sequencing result showing T-to-A mutagenesis of the target T (labelled red) in the 5′-TYTN-3′ DarT2 motif. 24 colonies were screened, out of which one contained the T-to-A mutation. The mutated bases are highlighted in red.

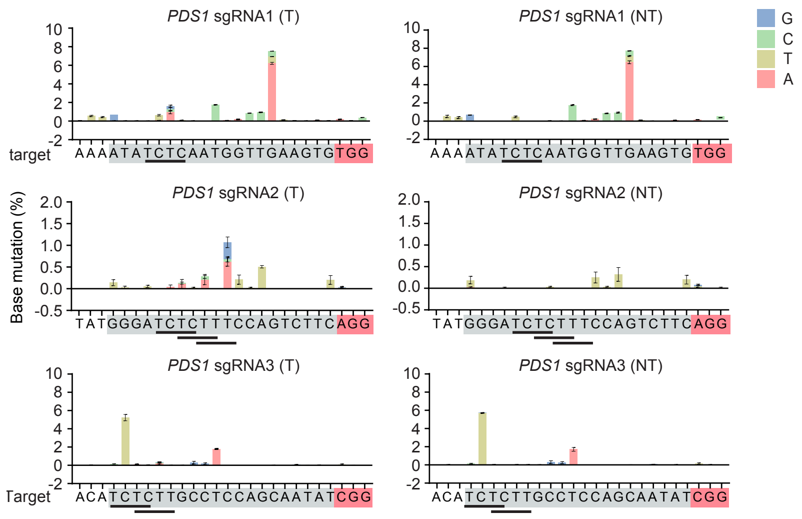

**Fig. S13. Location of base mutations with DarT2$^{D}$-SpnCas9 in the *PDS1* gene in *N. benthamiana* under targeting and non-targeting conditions.** Bars and error bars represent the mean and SEM of three independent replicates without selection or sorting. Peaks observed with both targeting (T) and non-targeting (NT) sgRNAs could represent amplification of *PDS1* homeologs. To account for these peaks, values from NT were subtracted from those from T to generate the plots shown in Figure 3g.

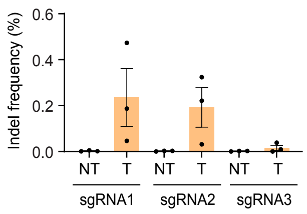

**Fig. S14. Frequency of indels generated with DarT2$^{D}$-SpnCas9 when targeting the *PDS1* gene of *N. benthamiana*.** Bars and error bars represent the mean and SEM of three independent biological replicates, where each replicate consists of tissue from four infiltrated leaves. Dots represent measurements from individual samples. T, targeting sgRNA. NT, non-targeting sgRNA.

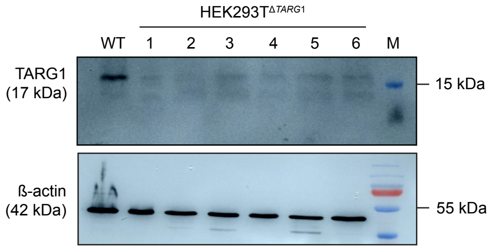

**Fig. S15. Confirming the *TARG1* knockout in HEK293T cell lines.** Clones 1-6 demonstrated disruption at the TARG1 targeted site using Sanger sequencing. Loss of protein expression was validated via western blotting using antibodies specific to human TARG1 (ref. [1]).

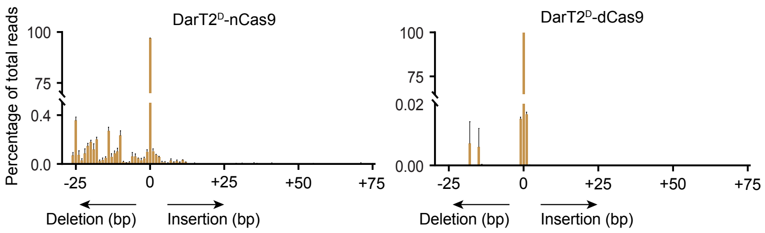

**Fig. S16. Size and frequency of indels when targeting the *EMX1* site 1 in HEK293T$^{\Delta TARG1}$ cells with DarT2$^{D}$-SpnCas9.** Values with '-' reflect size of deletions, '+' reflect size of insertions, while '0' represents the percentage of reads containing no insertions or deletions. Results derived from analysis of samples reported in **Fig. 4d.** Bars and error bars represent the mean and SEM of three independent replicates without selection or sorting.

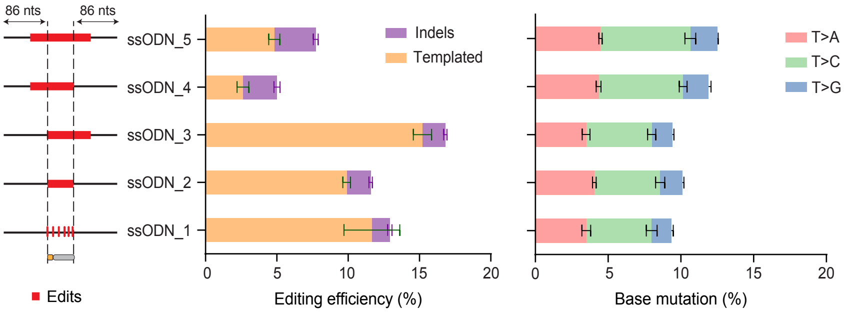

**Fig. S17. Base mutation with DarT2$^{D}$-SpnCas9 in HEK293T$^{\Delta TARG1}$ cells using different single-stranded oligodeoxynucleotides (ssODN) repair templates.** The editing assay was performed similar to **Fig. 4c**. Bars and error bars represent the mean and SEM of three independent replicates without selection or sorting.

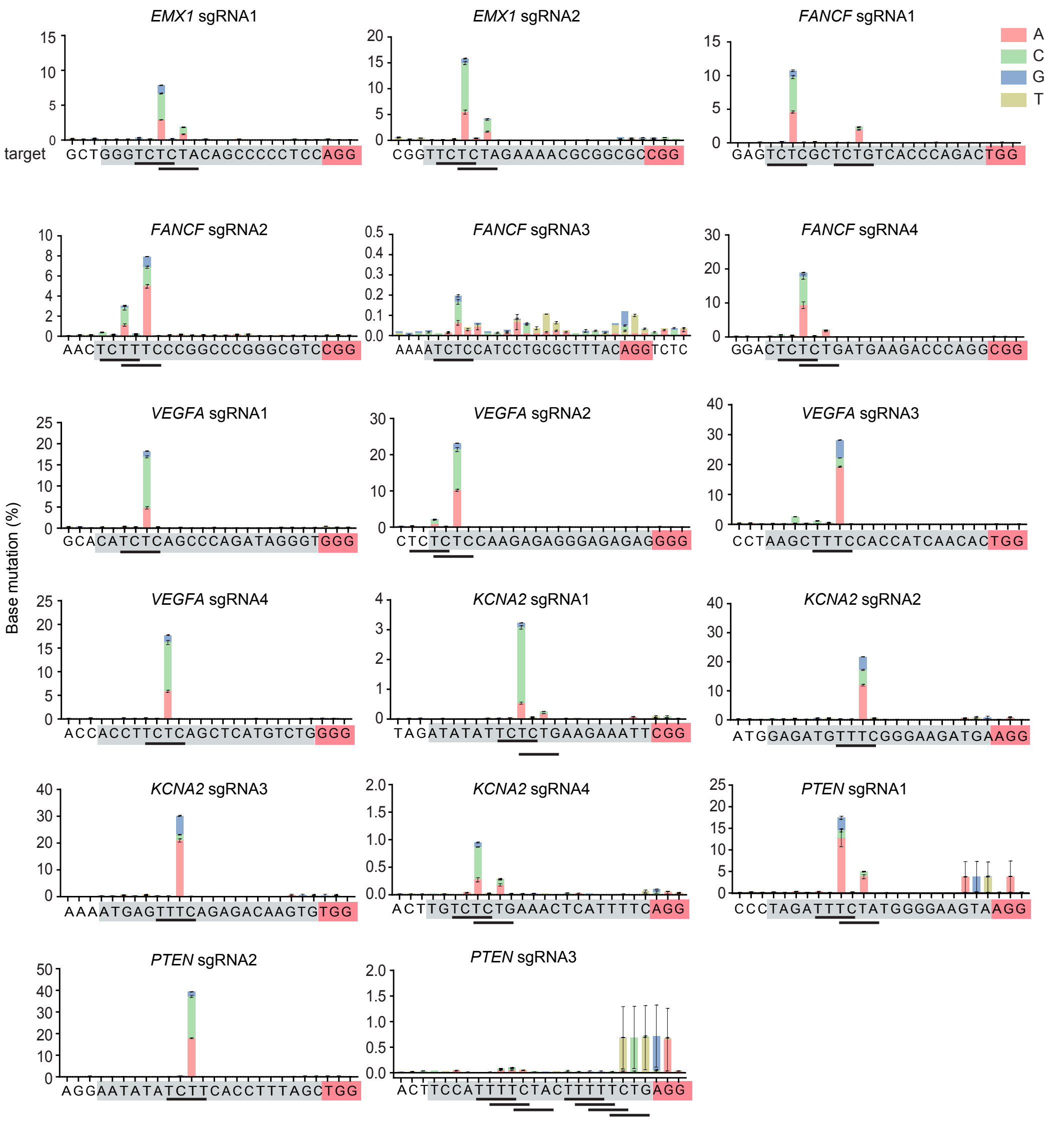

**Fig. S18. Location of base mutations with DarT2[D]-SpnCas9 in HEK293T-Δ*TARG1* cells.** Bars and error bars represent the mean and SEM of three independent replicates without selection or sorting. Horizontal bars indicate DarT2 recognition motifs, with ADP-ribosylation of the thymine at the third position.

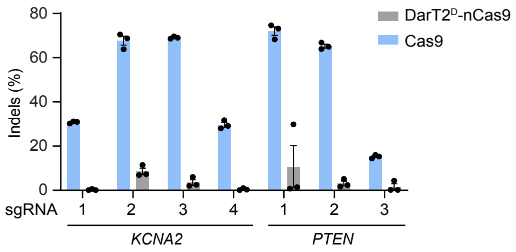

**Fig. S19. Quantification of total indels generated by DarT2(G49D)-nCas9 compared with traditional Cas9 in HEK293T$^{\Delta TARG1}$ cells.** Bars and error bars represent the mean and SEM of three independent replicates without selection or sorting. Dots represent individual measurements.

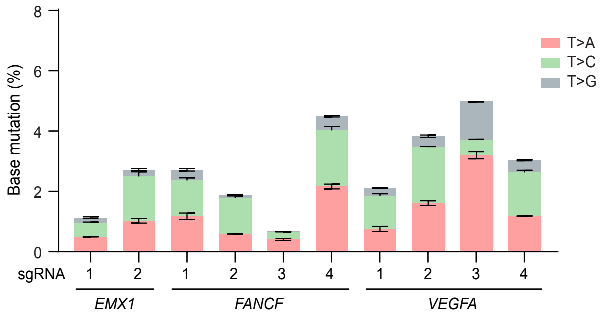

**Fig. S20. Base mutation frequencies at the ADP-ribosylated thymine with DarT2$^{D}$-SpnCas9 across different targets in *EMX1*, *FANCF* and *VEGFA* genes in U2OS$^{\Delta TARG1}$ cells.** Bars and error bars represent the mean and SEM of three independent replicates without selection or sorting.

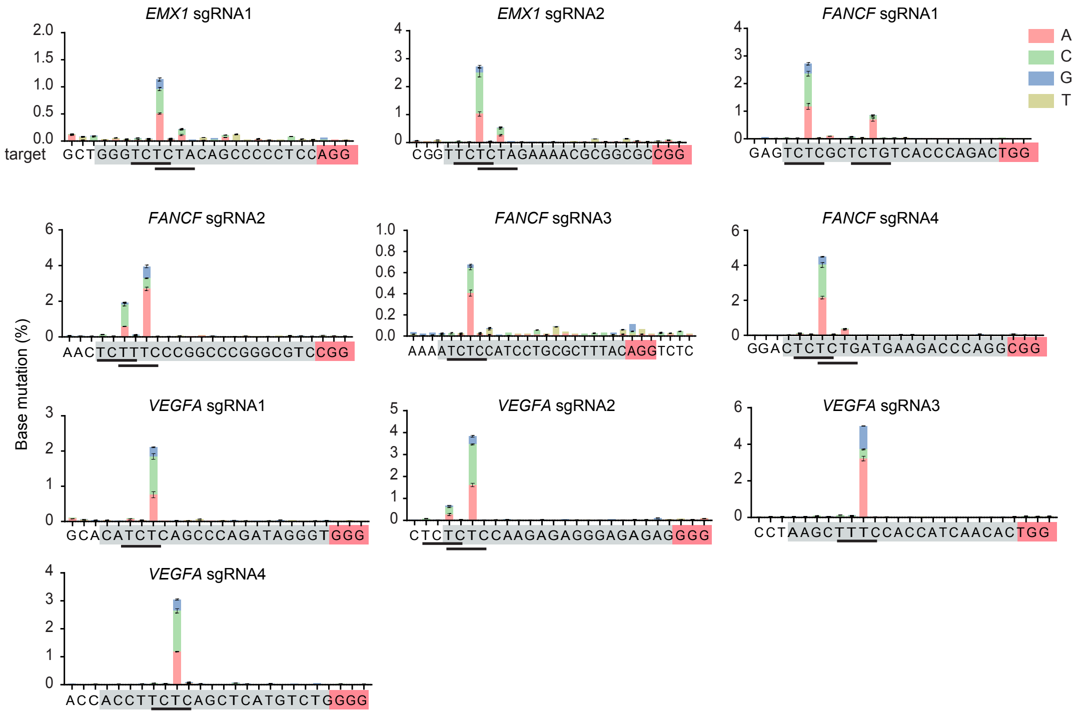

**Fig. S21. Location of base mutations with DarT2$^{D}$-SpnCas9 in U2OS$^{\Delta TARG1}$ cells.** Bars and error bars represent the mean and SEM of three independent replicates without selection or sorting. Horizontal bars indicate DarT2 recognition motifs, with ADP-ribosylation of the thymine at the third position.

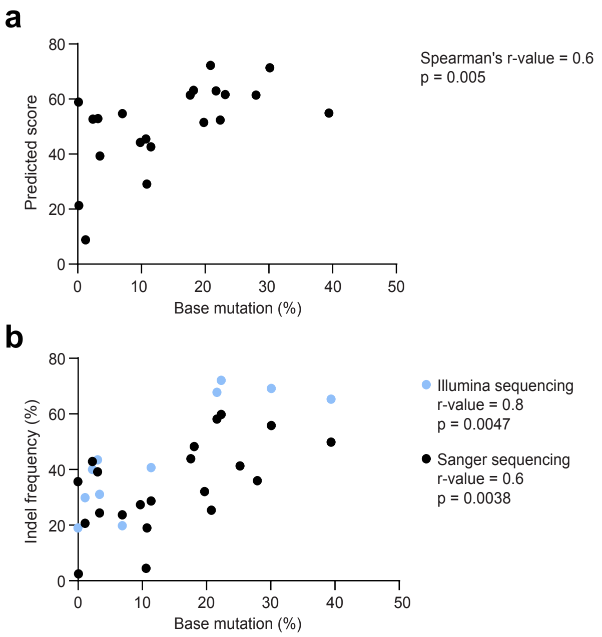

**Fig. S22. Correlation between measured or predicted indel formation at target sites with SpCas9 and ADPr-TAE-mediated base mutation. a,** Correlation between ADPr-TAE-mediated base mutation frequency and on-targeting scores predicted for indel formation with traditional Cas9. Each dot represents the mean of base mutation frequencies (n = 3) observed experimentally, plotted against scores predicted through the Rule Set 2 scoring method[2], for a specific genomic target. **b,** Correlation between ADPr-TAE base mutation frequency and indels generated by SpCas9. Each dot represents the mean of base mutation frequencies (n = 3), determined by Illumina sequencing against Indels determined through Illumina and Sanger sequencing methods (n = 3), for a specific genomic target.

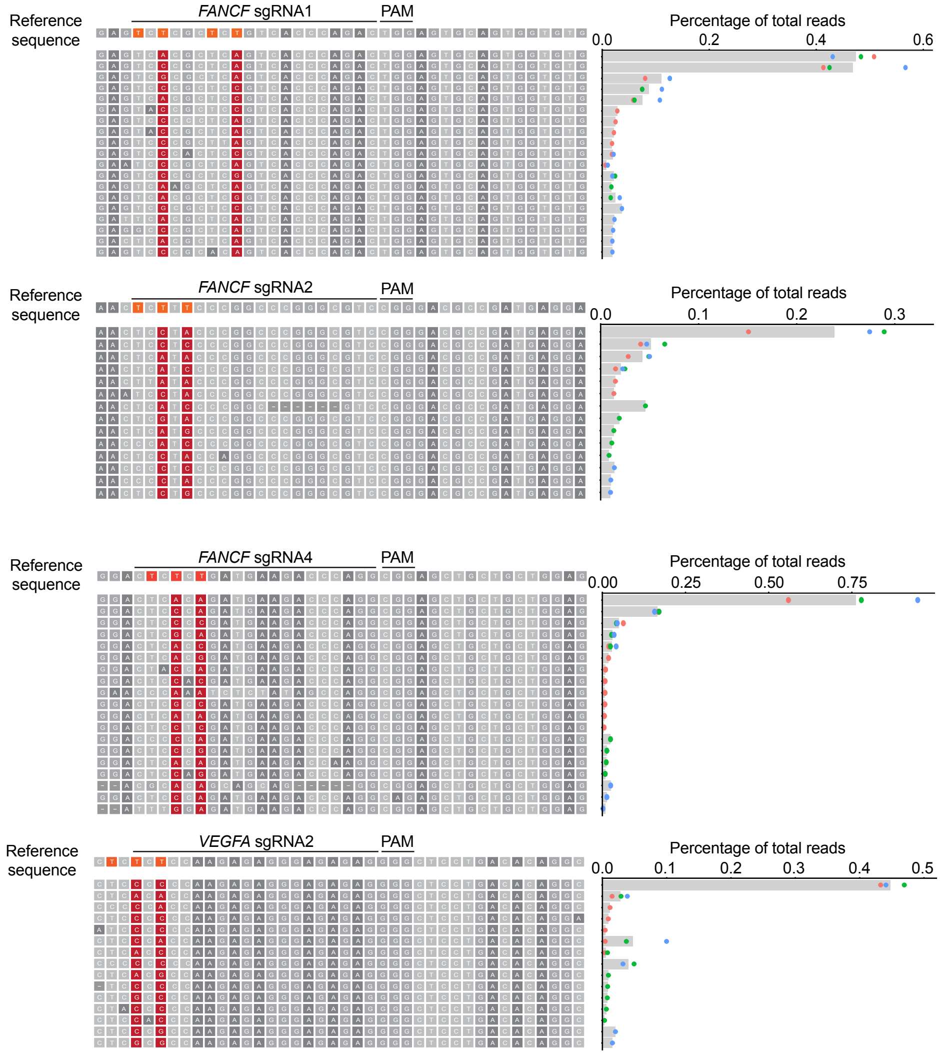

**Fig. S23. Frequency of co-occurring by-stander mutations when multiple motifs are present in the target sequence.** Data analysed from targets *FANCF*_sg1, *FANCF*_sg2, *FANCF*_sg4 and *VEGFA*_sg2 in **Fig. 4e** and **Fig. S7**. Values reported represent the mean and SEM of three independent replicates without selection or sorting. Dots represent measurements from individual replicates.

### SUPPLEMENTARY TABLES

**Table S1. Genome wide profiling of off-target single-nucleotide variations (SNVs) detected in the *kanR** strain under non-targeting conditions.** Values correspond to Figure 1i. SNVs were re-coded to the strand containing an edited C or T. Position three in the motif corresponds to the Genomic Position at which the SNV was detected. Genomic positions correspond to *E. coli* MG1655 (GenBank: U00096.3).

| Editor | SNV | Motif | Colony | Genomic Position | Read Depth | Percent |
|---|---|---|---|---|---|---|
| nCas9 | C>G | GTCG | 2 | 2884186 | 41 | 32 |
| nCas9 | C>G | TCCA | 3 | 2496515 | 40 | 28 |
| nCas9 | C>G | GTCG | 3 | 2884186 | 26 | 27 |
| nCas9 | C>G | CTCA | 3 | 4002022 | 37 | 35 |
| nCas9 | C>T | ATCG | 1 | 2238816 | 75 | 39 |
| nCas9 | T>C | ATTG | 1 | 1728463 | 76 | 28 |
| nCas9 | T>C | AGTG | 1 | 4032832 | 93 | 26 |
| nCas9 | T>C | GCTG | 2 | 1964854 | 83 | 27 |
| nCas9 | T>C | ATTG | 3 | 1996218 | 48 | 25 |
| DarT(G49D)-nCas9 | C>G | TCCA | 1 | 2496515 | 57 | 26 |
| DarT(G49D)-nCas9 | C>G | CTCA | 2 | 4002022 | 33 | 27 |
| DarT(G49D)-nCas9 | C>G | GTCG | 3 | 2884186 | 29 | 31 |
| DarT(G49D)-nCas9 | C>T | ATCG | 1 | 2238816 | 81 | 37 |
| DarT(G49D)-nCas9 | T>A | TTTC | 1 | 1741177 | 71 | 55 |
| DarT(G49D)-nCas9 | T>C | TTTC | 1 | 522283 | 58 | 48 |
| DarT(G49D)-nCas9 | T>C | TGTG | 2 | 4413038 | 52 | 25 |
| APOBEC-nCas9-UGI | C>T | CTCT | 1 | 1632710 | 58 | 45 |
| APOBEC-nCas9-UGI | C>T | CTCT | 1 | 2555285 | 137 | 47 |
| APOBEC-nCas9-UGI | C>T | CTCC | 1 | 2557310 | 127 | 46 |
| APOBEC-nCas9-UGI | C>T | CTCC | 2 | 237578 | 221 | 33 |
| APOBEC-nCas9-UGI | C>T | GTCC | 2 | 250724 | 461 | 37 |
| APOBEC-nCas9-UGI | C>T | TTCC | 2 | 373332 | 508 | 53 |
| APOBEC-nCas9-UGI | C>T | GTCA | 2 | 437138 | 513 | 51 |
| APOBEC-nCas9-UGI | C>T | GTCG | 2 | 1602770 | 464 | 32 |
| APOBEC-nCas9-UGI | C>T | TTCA | 2 | 2737123 | 484 | 56 |
| APOBEC-nCas9-UGI | C>T | CGCC | 2 | 3543825 | 498 | 98 |

| | | | | | | |
|---|---|---|---|---|---|---|
| APOBEC-nCas9-UGI | C>T | ATCG | 2 | 3561968 | 484 | 29 |
| APOBEC-nCas9-UGI | C>T | TTCC | 3 | 568237 | 251 | 33 |
| APOBEC-nCas9-UGI | C>T | TTCC | 3 | 569581 | 199 | 34 |
| APOBEC-nCas9-UGI | C>T | TTCC | 3 | 2189448 | 338 | 44 |
| APOBEC-nCas9-UGI | T>C | TGTG | 1 | 730239 | 77 | 26 |
| APOBEC-nCas9-UGI | T>C | ATTG | 1 | 3904655 | 116 | 29 |
| APOBEC-nCas9-UGI | T>C | GCTG | 1 | 4243748 | 124 | 26 |
| APOBEC-nCas9-UGI | T>C | AATG | 2 | 505166 | 483 | 25 |
| APOBEC-nCas9-UGI | T>C | AATG | 3 | 505166 | 440 | 25 |

**Table S2. Strains, plasmids, oligos and guideRNA spacers used in this work.** See the included Excel file.

**Table S3. Source data for Figures 1-4.** See the included Excel file.

**Table S4. Source data for Figures S1-S23.** See the included Excel file.

**Table S5. Potential disease targets for ADPr-TAE.** See the included Excel file.
